# Atlantic salmon farms are a likely source of *Tenacibaculum maritimum* infection in migratory Fraser River sockeye salmon

**DOI:** 10.1101/2021.06.15.448581

**Authors:** Andrew W. Bateman, Amy K. Teffer, Arthur Bass, Tobi Ming, Brian P. V. Hunt, Martin Krkošek, Kristina M. Miller

## Abstract

Infectious disease from domestic hosts, held for agriculture, can impact wild species that migrate in close proximity, potentially reversing selective advantages afforded by migration. For sockeye salmon in British Columbia, Canada, juveniles migrate past numerous Atlantic salmon farms from which they may acquire a number of infectious agents. We analyse patterns of molecular detection in juvenile sockeye salmon for one bacterial pathogen, *Tenacibaculum maritimum*, known to cause disease in fish species around the globe and to cause mouthrot disease in farmed Atlantic salmon in BC. Our data show a clear peak in *T. maritimum* detections in the Discovery Islands region of BC, where sockeye migrate close to salmon farms. Using well established differential-equation models to describe sockeye migration and *T. maritimum* infection spread, we fit models to our detection data to assess support for multiple hypotheses describing farm- and background-origin infection. Despite a data-constrained inability to resolve certain epidemiological features of the system, such as the relative roles of post infection mortality and recovery, our models clearly support the role of Discovery-Islands salmon farms in producing the observed patterns. Our best models (with 99.8% empirical model support) describe relatively constant (background) infection pressure, except around Discovery-Islands salmon farms, where farm-origin infection pressure peaked at 12.7 (approximate 95% CI: 4.5 to 31) times background levels. Given the evidence for farm-origin transfer of *T. maritimum* to Fraser-River sockeye salmon, the severity of associated disease in related species, and the imperilled nature of Fraser River sockeye generally, our results suggest the need for a more precautionary approach to managing farm/wild interactions in sockeye salmon.

## Introduction

Across taxa, long-distance migrations likely evolved to exploit resources, avoid predation, and – in some species – evade infectious disease (Altizer et al. 2011). Recent human agricultural practices, however, can result in elevated levels of infectious agents for migratory species passing or residing nearby (Krkosek 2010; Fritzsche McKay and Hoye 2016). If left unchecked, increasing agricultural incursions into previously undeveloped spaces may lead to elevated infectious-disease transmission at the wildlife-livestock interface, where human agriculture and wild species meet (Bengis et al. 2002; Wiethoelter et al. 2015). For cases in which long-distance animal migrations intersect the wildlife-livestock interface, associated infectious diseases may place a burden on migratory species, reducing or reversing the selective advantage of migration, whether or not it originally evolved to evade infection.

Globally, farmed and wild salmon (*Salmo* spp. and *Oncorhynchus* spp.) provide a well-studied example of the wildlife-livestock interface impacting migratory species. Although most likely not in response to disease risk, salmon have evolved to migrate long distances between freshwater spawning grounds and marine feeding grounds (Maekawa and Nakano 2002). Many populations now migrate past salmon farms near points of marine entry or exit (as juveniles and adults, respectively). Farmed salmon commonly share infectious diseases with wild salmon (compare: Nekouei et al. 2018; Thakur et al. 2018; Laurin et al. 2019), and active farms may augment natural host populations to push disease dynamics over critical density thresholds and generate outbreaks (Frazer et al. 2012; Krkošek 2017). Coastal salmon farms have been implicated in disease transfer to wild populations as well as associated population declines (Ford and Myers 2008; Costello 2009). Many relevant studies have focused on the impacts of parasitic sea lice (Krkošek et al. 2011; Frazer et al. 2012; Vollset et al. 2016), and while viral and bacterial diseases remain less explored, they have long been sources concern (Hastein and Lindstad 1991; Taranger et al. 2015).

In British Columbia, one high-profile example in which livestock may impact migration is that of sockeye salmon (*O. nerka*) and salmon farms. Multiple populations of Fraser-River sockeye salmon spawn in inland lakes and rivers, from which juveniles migrate downstream, past open-net salmon farms in the nearshore marine environment, and then on to feeding grounds in the northeast Pacific Ocean (Groot et al. 1991; Connors et al. 2012). Approximately 90% of Fraser River sockeye migrate north, between Vancouver Island and the mainland of BC (Welch et al. 2011), past about half of BC’s salmon farms. While time spent in the vicinity of individual farms is low (Rechisky et al. 2021), approximately 30 active farms, each farm holding 500 000 to 1 000 000 fish, lie along the main sockeye migration route between the Fraser River and the Alaska border in any given year (DFO, Figure 1). These farms can elevate the concentration of infectious agents faced by migrating juvenile sockeye (Shea et al. 2020), and depending on agent, elevated infectivity can extend for tens of kilometres beyond the confines of a farm (Krkošek et al. 2005; Foreman et al. 2015).

**Figure 1.**
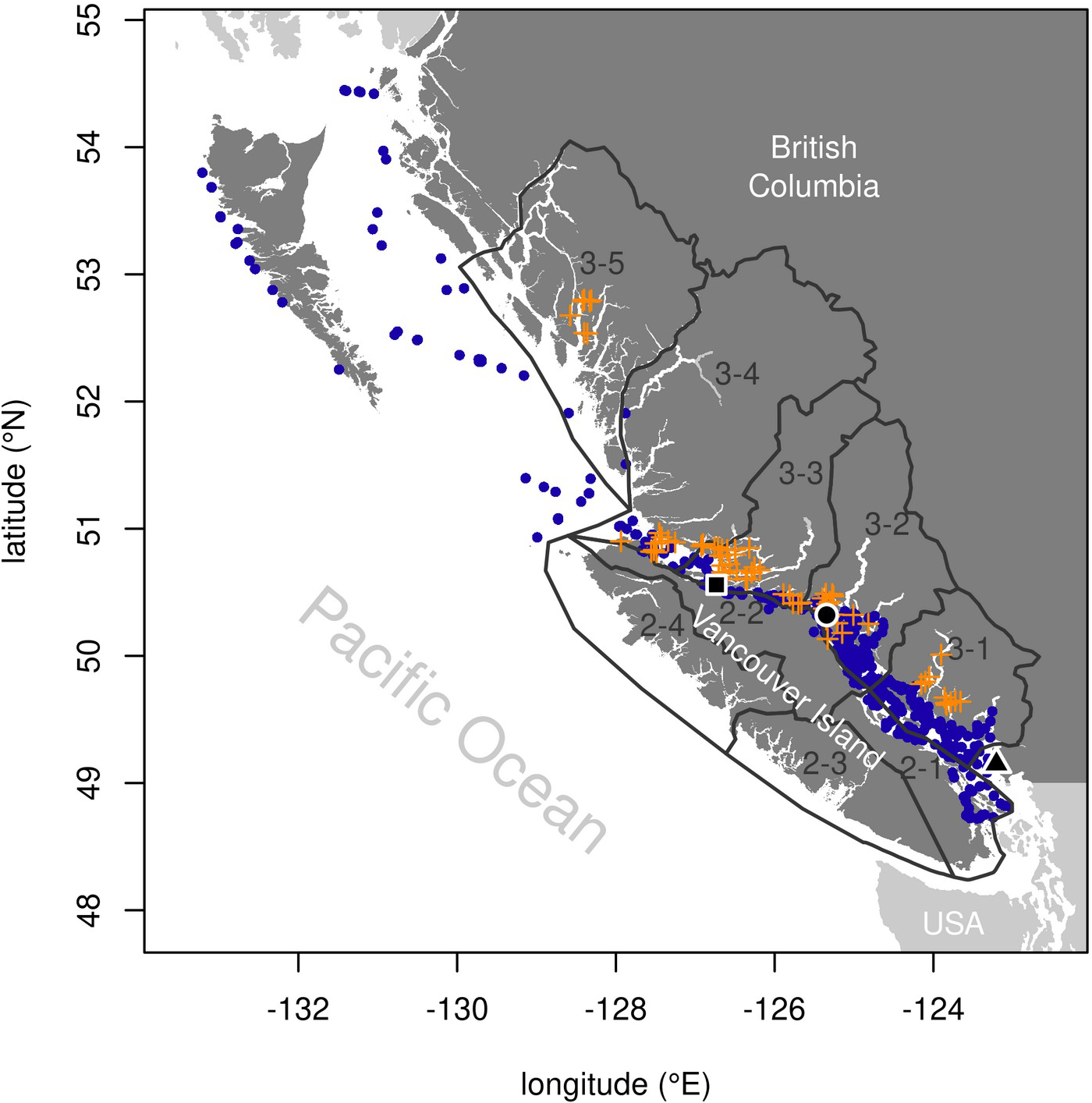
Collection locations for sockeye salmon (blue dots), along their north-westward migration after marine entry at the Fraser River mouth (black triangle), from which molecular detections of *Tenacibaculum maritimum* were analysed. Grey, labelled outlines indicate DFO’s Aquaculture Management Zones, and orange crosses show the locations of relevant open-net salmon farms (within zones 3-1 through 3-5). Black circle is centred on Discovery Islands, and black square indicates northern Johnston Strait.

Starting in about 1990, the number of maturing offspring per spawning adult (“productivity”) in Fraser-River sockeye declined, until a record-low number of spawners returning in 2009 sparked the Cohen Commission of Inquiry into the Decline of Sockeye Salmon in the Fraser River (Cohen 2012). At a cost of over $37 million (CAD), the Cohen Commission called scores of witnesses, reviewed thousands of pieces of evidence, and ultimately made 75 recommendations to promote the recovery of Fraser-River sockeye. The Commission cast particular scrutiny on the potential for disease transfer from salmon farms in the Discovery-Islands region of BC (Figure 1), where the majority of Fraser-River sockeye must migrate in close proximity to farms. A genomic signature consistent with viral disease was found to be associated with poor survival of Fraser sockeye (Miller et al. 2011), and analysis of productivity trends showed a link between Discovery-Islands farm production levels and Fraser-River sockeye productivity (Connors et al. 2012). A possible explanation for those trends is the transmission of infectious disease or other negative ecological interactions between farmed Atlantic salmon and wild sockeye salmon. Justice Cohen recommended that salmon farms in the Discovery Islands be removed if they were shown to pose any more than a minimal risk of serious harm to Fraser-River sockeye (Cohen 2012). In 2020, however, assessments by the Canadian federal department of Fisheries and Oceans (DFO) concluded that nine infectious agents, considered individually, each posed a minimal risk to Fraser-River sockeye as a result of potential transmission from salmon farms in the Discovery Islands (https://www.dfo-mpo.gc.ca/cohen/iles-discovery-islands-eng.html).

New evidence shows that one of the agents DFO assessed, *Tenacibaculum maritimum*, warrants further attention. A cosmopolitan marine bacterium that infects fish, *T. maritimum* infection does not invariably cause disease but is responsible for tenacibaculosis globally and causes “mouth rot” in Atlantic salmon (S. *salar*) on farms in BC (Frisch et al. 2018*b*). In a recent environmental-DNA (eDNA) study, out of 39 salmon pathogens for which presence was assessed in coastal marine waters, *T. maritimum* was one of the agents most strongly associated with active salmon farms (Shea et al. 2020). Here, we analyse new data from migrating sockeye salmon that were not included in DFO’s risk assessment (DFO 2020).

Using data pertaining to *T. maritimum* in juvenile sockeye-salmon smolts originating from the Fraser River, we developed and fitted a set of empirical spatio-epidemiological models to explore transmission-dynamic scenarios during the smolts’ early marine migration, from approximately April through August. Over the course of their migration, sockeye interact with many other populations of fish in addition to farmed Atlantic salmon, including other salmon species (congenerics) and Pacific herring (*Clupea pallasii*). We evaluate support for alternative hypotheses regarding farm- and background-origin infection to predict *T. maritimum* detection prevalence in sockeye smolts.

## Methods

We used sockeye-salmon samples and data from DFO and collaborators working on Juvenile Fraser River sockeye salmon in their first year of marine residence, caught during the course of existing marine sampling programs and sub-sampled for pathogen and molecular analysis. Specifically, sockeye came from DFO’s trawl programs and from the Hakai Institute’s purse-seine sampling program. Our analysis includes samples collected in the spring and summer from 2008 through 2018. In order to evaluate evidence for infection from salmon-farm versus background sources between Vancouver Island and the mainland of BC, we restricted our analyses to fish that were caught between Vancouver Island and the mainland of BC and further along the sockeye migration route to the northwest (Figure 1), and we used genetic stock ID (Beacham et al. 2004, 2010) to further restrict our analysis to sockeye smolts from stocks known to leave the Fraser River and primarily migrate north through that region (Table S1).

We screened the selected sockeye for the presence of *T. maritimum* (among other infective agents) using molecular techniques described in detail elsewhere (Fringuelli et al. 2012; Miller et al. 2016). Briefly, the methods involve nucleic acid extraction followed by quantitative polymerase chain reaction (qPCR) screening on the Fluidigm BioMark™ nanofluidics platform. This approach estimates specific nucleic-acid-sequence copy number within a fixed-volume sample at standardised nucleic-acid concentration via comparison to known-concentration serial dilutions of artificial construct standards of the targeted sequence. For the purposes of our models, we reduced *T. maritimum* nucleic acid copy number data to a per-fish measure of detection. If duplicate qPCR tests on the same sample both detected the bacterium, we considered that sample to be positive; if both tests indicated a lack of the bacterium we considered the sample negative; and if the two tests disagreed we considered the sample indeterminate and excluded it from further analyses. Resulting molecular *T. maritimum* detection data were available for 2270 Fraser-River sockeye smolts.

To make the sockeye-migration tractable, we simplified the two-dimensional sampling locations into measures of migration distance along a single spatial axis, representing least-seaway distance (generally to the northwest) from the Fraser River mouth. To do this, we considered all over-water straight-line paths among sample-collection and shoreline points, and then solved for the shortest paths between sampling locations and the Fraser River mouth using Dijkstra’s algorithm (Dijkstra 1959; Csardi and Nepusz 2006). We assigned negative migration values to sampling locations southeast of the Fraser River.

Detection data from sockeye-smolt screening showed a distinct peak in *T. maritimum* prevalence near salmon farms in the Discovery Islands region of BC (Figure 2). The vast majority of samples collected before and after the Discovery Islands, from the perspective of a migrating sockeye smolt, yielded no *T. maritimum* detections. Despite inevitable variability, samples from the vicinity of the Discovery Islands showed a pattern of substantially higher detection prevalences.

**Figure 2.**
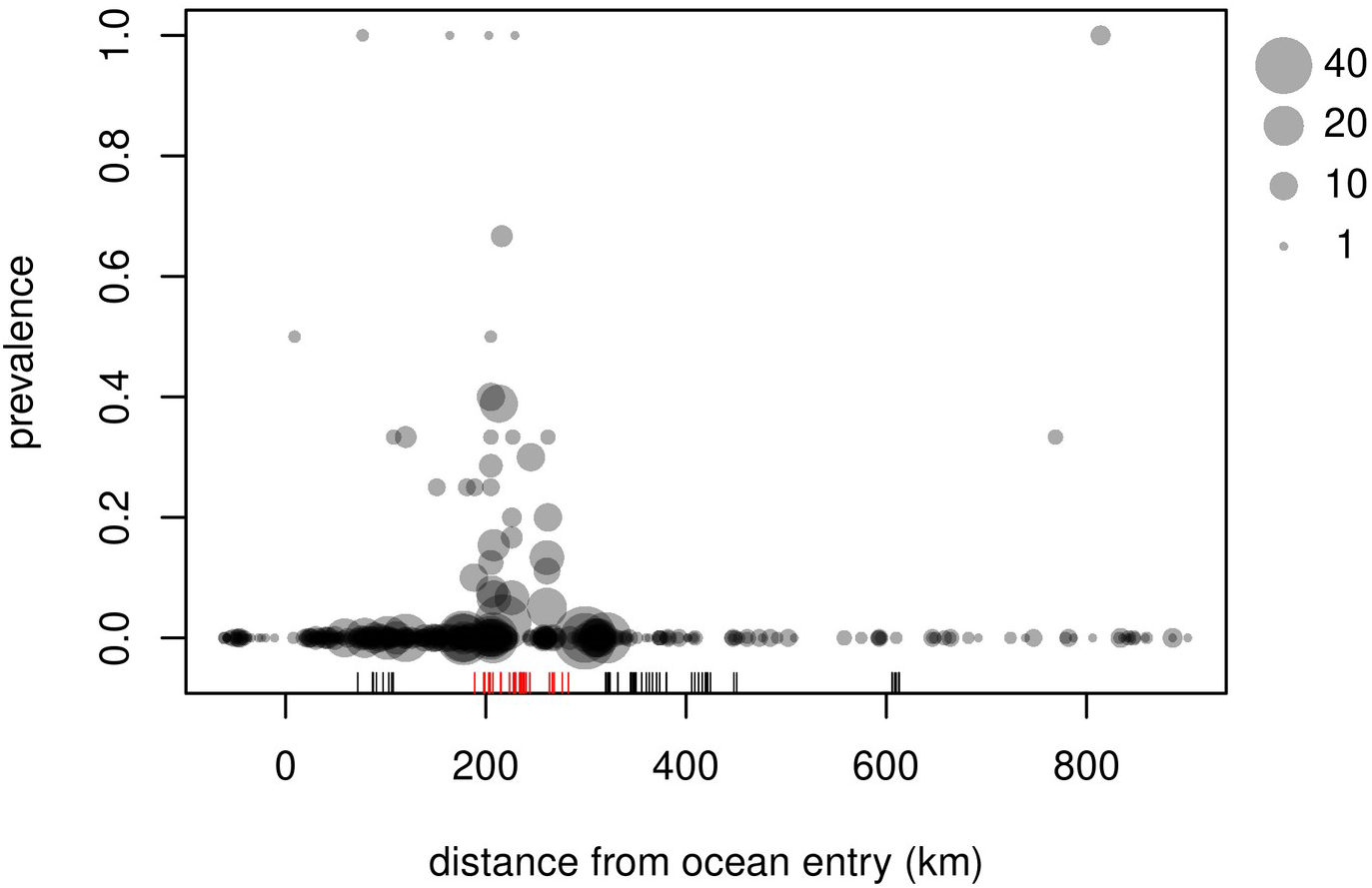
Prevalence of *Tenacibaculum maritimum* infection, detected via high-throughput qPCR assay, in samples of Fraser-River sockeye salmon smolts migrating north-westward from the mouth of the Fraser River, BC (*x*=0). Grey circles represent smolts caught in individual trawl and purse-seine surveys between 2008 and 2018. “Rug” shows locations of salmon farm tenures, with red indicating farms in the Discovery Islands region of BC.

Although it might seem natural to interpret these data as evidence that infections originating in the Discovery Islands, and the salmon farms located there, largely determine *T. maritimum* infection patterns in Fraser-River sockeye salmon, we must consider several possible alternative explanations. First, any infection can take time to develop, and the spatial *T. maritimum* infection pattern may be a consequence of detections that are spatially displaced from their precipitant exposure locations, even if infection mainly comes from largely constant background sources. Second, the pattern of infection may be best described by spatially varying background source pressure, not directly tied to salmon farms and perhaps due to idiosyncrasies of local hydrodynamic patterns (Chandler et al. 2017; Khangaonkar et al. 2017). The Discovery Islands region serves as a hydrographic funnel that concentrates all out-migrating salmon into a confined space where interactions with background infection sources may be amplified. Third, even if salmon farms are a major source of *T. maritimum* infection for Fraser-River sockeye, farms in the Discovery Islands specifically may or may not provide an outsized contribution to infection pressure. In this latter case, the Discovery-Islands region may be where we happen to see elevated infection rates, again due to a combination of migration and infection processes.

Our detailed analyses, described below, confronted these hypotheses, formalised as mathematical models, with the data at hand. By comparing empirical support for different models, we show which hypotheses are most consistent with observed patterns.

### Model overview

We assembled models from well-established components to describe infection dynamics for Fraser-River sockeye smolts as they migrated away from the mouth of the Fraser River. The partial differential equation (PDE) models we used combined two basic parts: a spatial component, to describe the changing spatial distribution of sockeye as the population migrates during a typical year, and an epidemiological component, to describe exposure to *T. maritimum*, development of associated infections, and subsequent recovery from or mortality due to those infections.

The spatial component of our models took the form of standard advection-diffusion equations (Murray 2007). This class of models tracks the abundance or density of individuals over space and time, in our case describing directed movement (advection) associated with migration, as well as spread (diffusion) due to variation in migration speed or milling behaviour. Technically, we modelled the spatial probability-density function for migrating sockeye that entered the marine environment in a given year. The models also capture natural mortality over the course of migration (a “decay” component), so that the spatial density of sockeye integrates to a smaller and smaller fraction over time, regardless of how that density is distributed.

The epidemiological component of our models took a standard “susceptible-exposed-infected-susceptible” (SEIS) form, whereby the susceptible portion of the population can become exposed, after which infection develops at a given rate. This model requires modification of the spatial model by decomposing the time-varying spatial distribution of sockeye into susceptible, exposed, and infected components. Use of this epidemiological description means that the model overall must be formulated as a set of advection-diffusion-reaction equations. We assume that sockeye leave freshwater as uninfected, susceptible individuals. Susceptible fish can become exposed due to spatially varying infection pressure in the marine environment, but we assume that molecular detection of *T. maritimum* is impossible until those exposed fish develop into infected fish. Infected fish may die at a faster rate and can recover and return to the susceptible category.

The overall model takes the form:

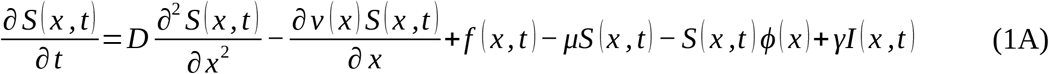

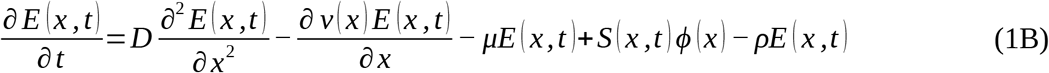

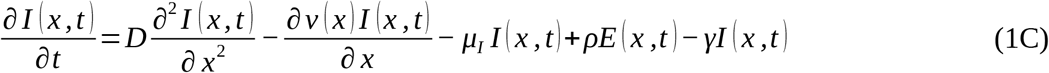

(For convenience, in much of the subsequent text, we drop the explicit space- and time-dependence.) Here, *S*, *E*, and *I* are the space- and time-varying relative densities of susceptible, exposed, and infected individuals; *x* is the single spatial migration-distance dimension; *t* is time; *D* is the spatial diffusion coefficient and *v* is the migration speed (below, we describe how *v* changes in space); *f* represents the time-varying input of sockeye smolts at the Fraser River mouth; *μ* is mortality (with a different rate, *μ_I_*, for infected individuals); *ϕ* represents the spatially varying cumulative infection pressure (below, we describe how this is modelled), *ρ* is the development rate of exposed into infected individuals; and *γ* is the recovery rate of infected individuals. This model form assumes that sockeye salmon can become infected with *T. maritimum* from external sources but, given the short time window during which they can be sampled, relevant secondary transmission does not occur.

### Movement model portion

To maximise our chances of being able to fit the overall model (Equation 1) to detection-prevalence data, we first parameterised the movement portion of the model (Equation S1) using additional information from a number of sources. One missing piece was an estimate for the diffusion coefficient, *D*. As a result, in the course of numerically solving the model (technical details below), we tuned *D* so that movement-model predictions aligned with observations of migration timing at two points along the Fraser-sockeye migration route.

Where information was available, we drew parameter estimates from the literature. We used the approximate long-term mean and standard deviation of sockeye-smolt captures from the Mission rotary screw trap (Preikshot et al. 2012) to parameterise a Gaussian form for the input function, *f*. This involved a mean Julian day for smolt passage at Mission of 124 (May 4^th^), plus a correction accounting for the distance from Mission to the Fraser River mouth of 75 km and a lower-Fraser swimming speed of 187 km/day (Stevenson et al. 2019), and a standard deviation of nine days (Preikshot et al. 2012). We used migration speeds, *v*, from a tracking study of age-one smolts (Stevenson et al. 2019), with values of 9.5 km/day from the Fraser River to the Discovery Islands region, and 14.5 km/day thereafter. We set the baseline mortality rate, *μ*, to 0.00924 such that survival over the course of 150 days would be 25%, to approximately align with Stevenson et al. (2019). Refitting our best model (see Results) with *μ* = 0.00611, to produce 40% survival after 150 days, as seen for larger but rare age-2 smolts (Bass et al. 2020), yielded near-identical results.

We numerically solved the susceptible-only advection-diffusion-decay model in R (R Core Team 2020) using the “method of lines,” as implemented in the *ReacTran* package (Soetaert and Meysman 2010). We used a 1 km spatial discretisation extending from *x* = −100 km (southeast of ocean entry) to *x* = 1500, and we used a 1 day temporal discretisation from Julian days 75 (March 15^th^) through 275 (October 2^nd^). Initial density, *S*(*x*,75), was zero, and we set zero-flux boundary conditions for *x*. We set the diffusion coefficient, *D*, by iteratively matching the modelled cumulative distribution of smolts that had passed the Discovery Islands (*x* = 211 km) and northern Johnstone Straight (*x* = 309 km; Figure 1) to corresponding cumulative distributions of catch-per-unit-effort from purse-seine sampling run by the Hakai Institute between 2015 and 2019 (Johnson et al. 2019). Resulting model predictions from the advection-diffusion-decay movement model matched catch data well (Figure 3).

**Figure 3.**
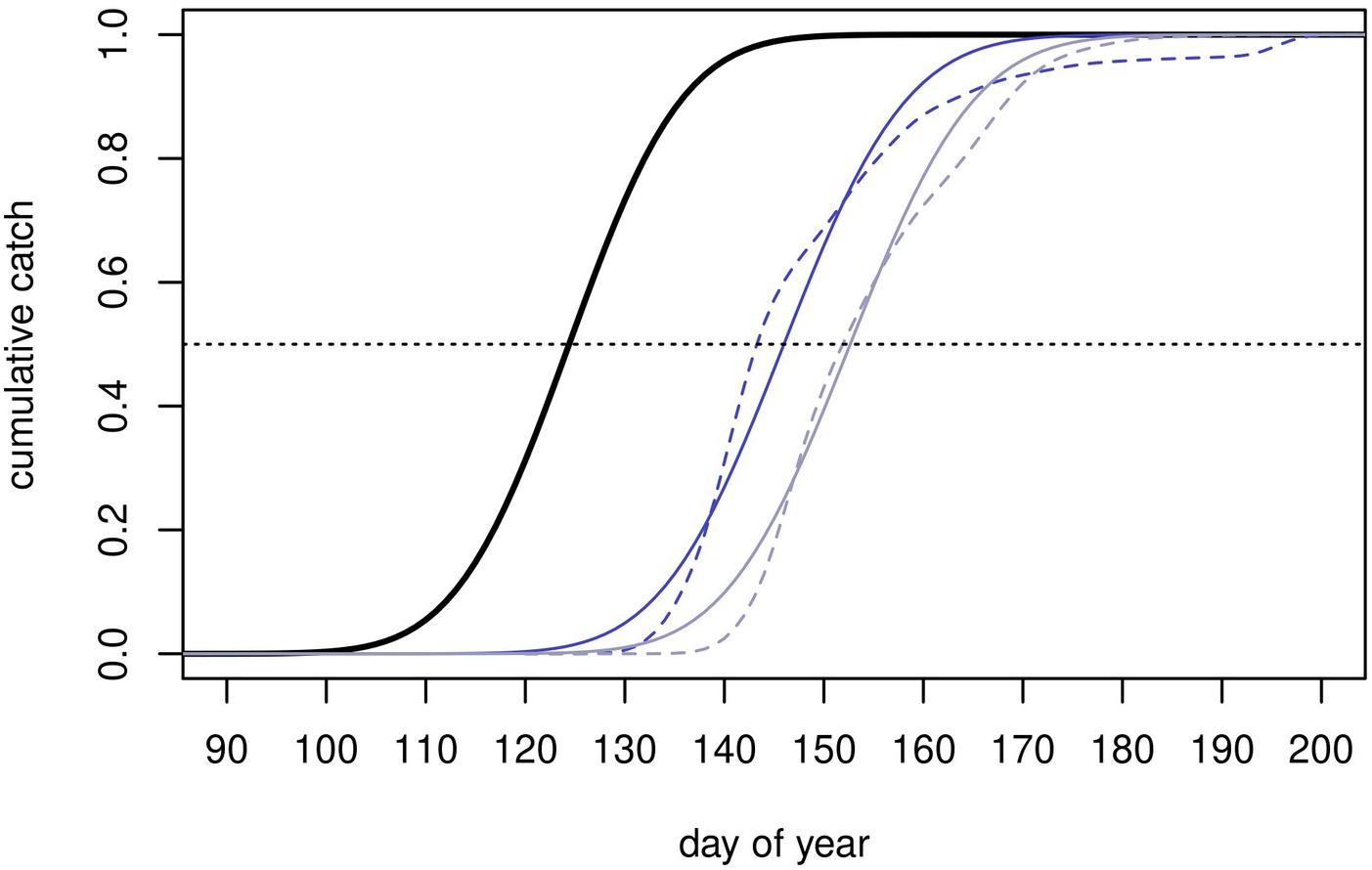
Cumulative Fraser-River sockeye salmon smolt passage at Mission, BC (black curve, parameterised from long-term data) and in the Discovery Islands (blue; 211 km from ocean entry) and northern Johnstone Straight (grey; 309 km from ocean entry) regions of their north-westward migration route. For the latter two locations, solid curves come from a parameterised advection-diffusion-decay model of sockeye movement, while dashed curves show empirical data from 2015 through 2019, collected by the Hakai Institute & colleagues.

### Epidemiological model portion

Arguably the most important component of the epidemiological portion of Equation (1) is the infection pressure, *ϕ*, critical for drawing inference about the source of *T. maritimum* infection in juvenile sockeye. To model *ϕ* we considered four candidate models based on the sum of putative infection pressures from background and salmon-farm sources (*ϕ_B_* and *ϕ_F_*, respectively); *ϕ* = *ϕ_B_* + *ϕ_F_*. Two candidate sub-models considered background sources only: a constant background infection pressure for all *x*, at level *β*_0_, to be determined at the time of model fitting; and background pressure that varied by DFO Fish Health Zone (Table 1), at levels *β*_1_, *β*_2_, *β*_3_, *β*_4_, and *β*_5_. These infection pressures were determined at the time of model fitting, via one and five free parameters, respectively. Two candidate sub-models considered farm-source infection in addition to constant background pressure: one accounted for farm activity and distance from farms, and the other modified the first, allowing farms in the Discovery Islands to contribute proportionately more or less to overall farm-source infection pressure.

**Table 1:** Migration-distance cutoffs used in spatio-epidemiological models to approximate DFO Aquaculture Management Fish Health Zones along the north-westward migration of juvenile Fraser-River sockeye salmon. Boundary locations were generated by comparing migration distance (*x*) estimates at sockeye sampling locations to known zonal boundaries.

| Fish Health Zone | southern boundary (km) | northern boundary (km) |
| --- | --- | --- |
| 3-1 & south | - | 130 |
| 3-2 | 130 | 234 |
| 3-3 | 234 | 343 |
| 3-4 | 343 | 410 |
| 3-5 & north | 410 | - |

The first farm-source infection sub-model considered infection risk that declined with distance from each active farm according to a Gaussian “dispersal” kernel, assuming an average source strength proportional to total months of farm activity in our study period. We used a Gaussian kernel as a coarse phenomenological description of bacterial dispersal around a farm, reasoning that the cumulative infection pressure from multiple farms would likely have more bearing than the kernel’s particular functional form. Scaling source strength to proportional farm activity (Salama and Murray 2011) effectively modelled infection pressure at a typical point in time, since we deemed that data constraints made a complex time-varying model of infection pressure infeasible for overall model fitting. The per-farm contribution to infection-pressure took the form:

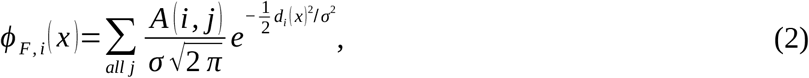

where *A*(*i,j*) is an indicator function that is one if farm *i* was active in month *j* and zero otherwise, *σ* is the standard deviation of the kernel to be determined at the time of model fitting, and *d_i_*(*x*) is a distance function for each farm. The *d_i_*(*x*)’s account for the facts that migrating sockeye approach and then pass each farm along their migration route but many farms do not lie directly on that route. We empirically approximated each farm’s *d_i_*(*x*) by fitting a shifted absolute-value function of migration distance (*x*) to seaway distances from farm *i* to the sampling location of each sampled fish (e.g. Figur*e S1*). Each *d_i_*(*x*) function had parameters for the minimum value, *a_i_*, and *x*-value at that minimum, *x_0,i_*, and was fit assuming normally distributed residuals with standard deviation *σ_d,i_*. The final version of the farm infection pressure was the scaled sum, across all farms, of the per-farm contribution in Equation (2). Further, although we simplified our modelling framework to consider only a single spatial axis, *x*, the real population of migrating fish is spread out perpendicular to our migration axis. To account for variation in the distance to each farm, as experienced across the population at each migration distance, we averaged across the residual variation from our *d_i_*(*x*) fits (i.e. the distribution of distance to farm *i* for fish caught at location *x*), so that:

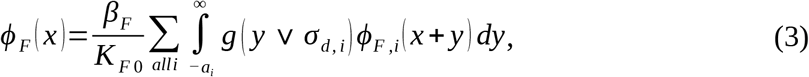

where *β_F_* is a scaling parameter also to be determined at the time of fitting, *K*_*F*0_ is a standardisation factor (the maximum value of the sum to the right, over all *x*), *g*(*y*|*σ_d,i_*) is the Gaussian probability density function with standard deviation *σ_d,i_* and mean 0 evaluated at *y*. The lower limit of the integral is set to avoid considering nonsensical negative farm-distance values at *x_0,i_*, where migrating individuals come closest to the farm and the highest associated infection pressures will occur. The *K*_*F*0_ factor ensures that *β_F_* is the maximum value of *ϕ_F_*, for ease of comparison to *ϕ_B_*. In practice, to facilitate repeated calculation during model fitting, we approximated *ϕ_F_* by replacing the integral in Equation (3) with a Reiman sum from −3*σ_d,i_* (or −*a_i_*, whichever was greater) to 3*σ_d,i_* with 0.1 km intervals. To manage numerical errors, we also imposed a lower bound of 0.1 km on *σ* (the standard deviation of the infection kernel) so that any infection kernel was not “skipped over” by too coarse a discretisation.

The second version of the farm-source infection sub-model allowed for different source strengths for Discovery-Island farms relative to other farms along the sockeye migration route. To achieve this, we replaced *A*(*i,j*) in Equation (2), where *i* corresponded to Discovery-Island farms, with *e^δ^A*(*i,j*). Here, *e^δ^* is a scaling factor, with *δ* > 0 leading to elevated (and *δ* < 0 leading to reduced) infection pressure from Discovery-Island farms.

For all versions of the epidemiological sub-model, we considered the development rate (*ρ* > *0*) and recovery rate (*γ* > *0*) to be free parameters, determined at the time of model fitting, and we modelled infected mortality as a scaled version of baseline mortality: *μ_I_ = e^η^μ*. Here, *e^η^* is an inflation factor (*η* > *0* assumed, also treated as a free parameter) that controls mortality in infected individuals, relative to background mortality.

### Model fitting and comparison

We treated the parameterised movement portion of the model as fixed, given the quality of data we used for parameterisation. We also assumed movement parameters to be the same for each epidemiological class of fish in Equation (1). This allowed us to focus model fitting on unknown parameters related to infection pressure(s) and progress.

In fitting the epidemiological components of the model to the *T. maritimum* detection data from sockeye smolts, we used the same approach to numerically solve Equation (1) as we had for the movement model (Equation S1) alone. Assuming that molecular detections were from infected fish only, we took the infection status of each fish, *k*, sampled at location *x*(*k*) and time *t*(*k*), to have come from a Bernoulli distribution with probability of infection:

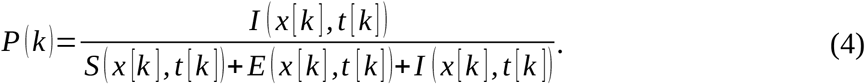

Thus, the likelihood for the infection status of fish *k* was 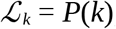, if *T. maritimum* was detected, and 1-*P*(*k*) otherwise. In fitting the model, we minimised the negative log of 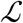, the joint likelihood across all screened sockeye salmon. Assuming independence among observations, this was:

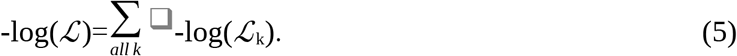

We minimised equation (5) with R’s *optim* numerical optimiser, using the Nelder-Mead algorithm to find the combination of model parameters that minimised each model’s negative log-likelihood, given the data. That is, we solved for the maximum-likelihood estimates (MLEs) of the parameters. For each of the four infection sub-models, we considered both a SEIS version of the PDE model (1) and a slightly less complex “SIS” version (Equation S2), in which exposed individuals immediately become infected, without passing through a separate exposed class. Overall, this meant that we fit eight candidate models: an SEIS and SIS version with each of four infection sub-models (Table 2).

**Table 2:**
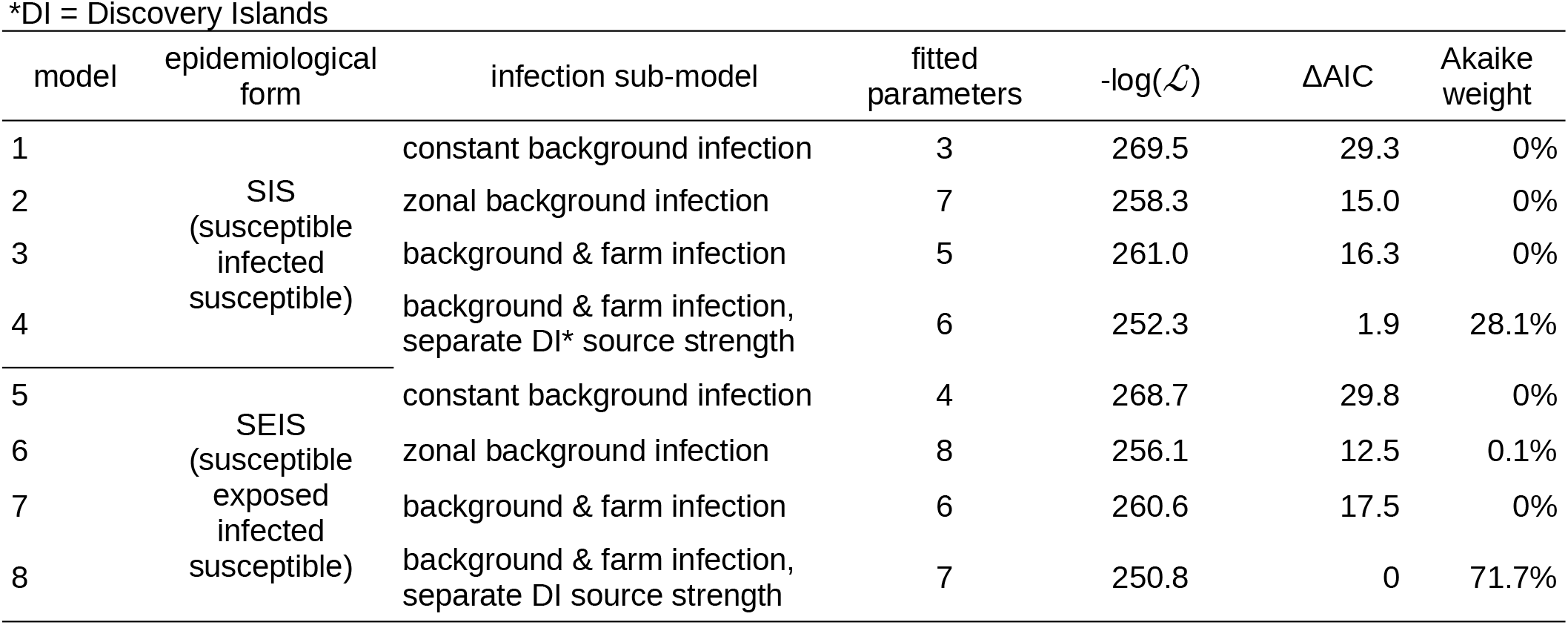
Comparison of fitted spatio-epidemiological candidate models to describe patterns of *Tenacibaculum maritimum* detection in Fraser River sockeye salmon smolts during their early marine migration.

For each model, we calculated Akaike’s information criterion (Akaike 1973) and ΔAIC scores (difference in AIC relative to the lowest-AIC model). Lower ΔAIC values indicate “better” (more parsimonious) models, with values below two indicating substantial model support, values above four indicating considerably less support, and values above ten indicating essentially no support (Burnham and Anderson 2002). Associated Akaike weights represent the strength of evidence for each model, given the candidate model set.

### Parameter and model uncertainty

We initially attempted to estimate parameter confidence intervals via the quadratic-approximation method (Bolker 2008). The scaling parameter for the Discovery-Island farm infection strength, *δ*, caused problems, however; the Hessian (second-derivative “curvature” matrix) of the negative log likelihood function, evaluated at the MLEs, could not be inverted to yield the approximate variance-covariance matrix for the parameter estimates (Bolker 2008). As a result, we used likelihood profiles, fixing one or two parameters at a sequence of values spanning their MLEs and optimising all other parameters to explore the likelihood function. We first generated a likelihood profile and associated confidence interval for the *δ* parameter, which indicated that large values of *δ* all produced an effectively equivalent model fit (Figure S2A). In light of this result, to improve the stability of the optimiser and allow estimation of a variance-covariance matrix for the majority of model parameters, we used a Discovery-Island-only version of the farm-infection sub-model, *ϕ_F_*, for further uncertainty calculations. That is, we set farm contributions outside of Fish Health Zone 3-2 (Table 1) to zero. Our resulting confidence estimates may thus be slightly too small (since a variable *δ* could improve the model fit at each combination of the other parameters); however, given the clear results with respect to *δ*, we deemed it useful to present uncertainty estimates, approximate though they are. Using the *δ*- restricted model, we generated a likelihood profile and associated confidence interval for σ, the parameter controlling the spread of infection around individual farms. We also generated joint likelihood profiles and associated confidence regions for: 1) the infection parameters (*β*_0_ and *β_F_*) and 2) the infected mortality (*μ_I_*) and recovery (*γ*) parameters.

To generate confidence regions for predictions from the best model (subject to the Discovery-Island-only modification to improve numerical stability), we drew 1000 samples from a multivariate normal distribution with means equal to the MLEs and variance-covariance matrix estimated from the curvature of the negative log likelihood at the MLEs. For each of these parameter combinations, we generated model predictions, and at each spatiotemporal location (*x,t*) we generated desired percentiles, representing the specified range of uncertainty associated with the fitted model.

### Inter-annual variation

We base inference on models fit to data from all years of our study. In part, this was due to a lack of comprehensive data (e.g. incomplete annual migration-timing data at Mission, inconsistent relative abundance measures along the sockeye migration route). Inter-annual variation in infection rates does, however, occur. In 2015, in particular, we saw high *T. maritimum* infection rates in Fraser-River sockeye smolts in the marine environment. To explore how this might have affected our conclusions, we refit the best version of the epidemiological sub-model using data from 2015 alone, and to data from all other study years. In both cases, *T. maritimum* detection data and farm-stocking data were specific to each time period.

## Results

The best partial-differential-equation model of *T. maritimum* infection prevalence in juvenile Fraser-River sockeye was the SEIS version in which farm-source infection differed between Discovery-Islands and other farms (Model 8; Table 2). This model accounted for 71.7% of model support, and when considered along with the corresponding SIS model (Model 4; Table 2), the associated version of the infection sub-model accounted for 99.8% of model support. The best model described a pattern of infection prevalence that was relatively constant throughout the sockeye migration route, except around Discovery-Island farms, where infection was elevated (Figure 4). While the movement model described a wave of sockeye population density travelling north (Figures 2, S3A), the spatial pattern of infection prevalence changed little over the course of the migration season (Figure S3B). These patterns combined to produce a peak in population-wide infection prevalence just before Julian day 150 (May 30^th^), as most sockeye passed the Discovery Islands (Figure 5).

**Figure 4.**
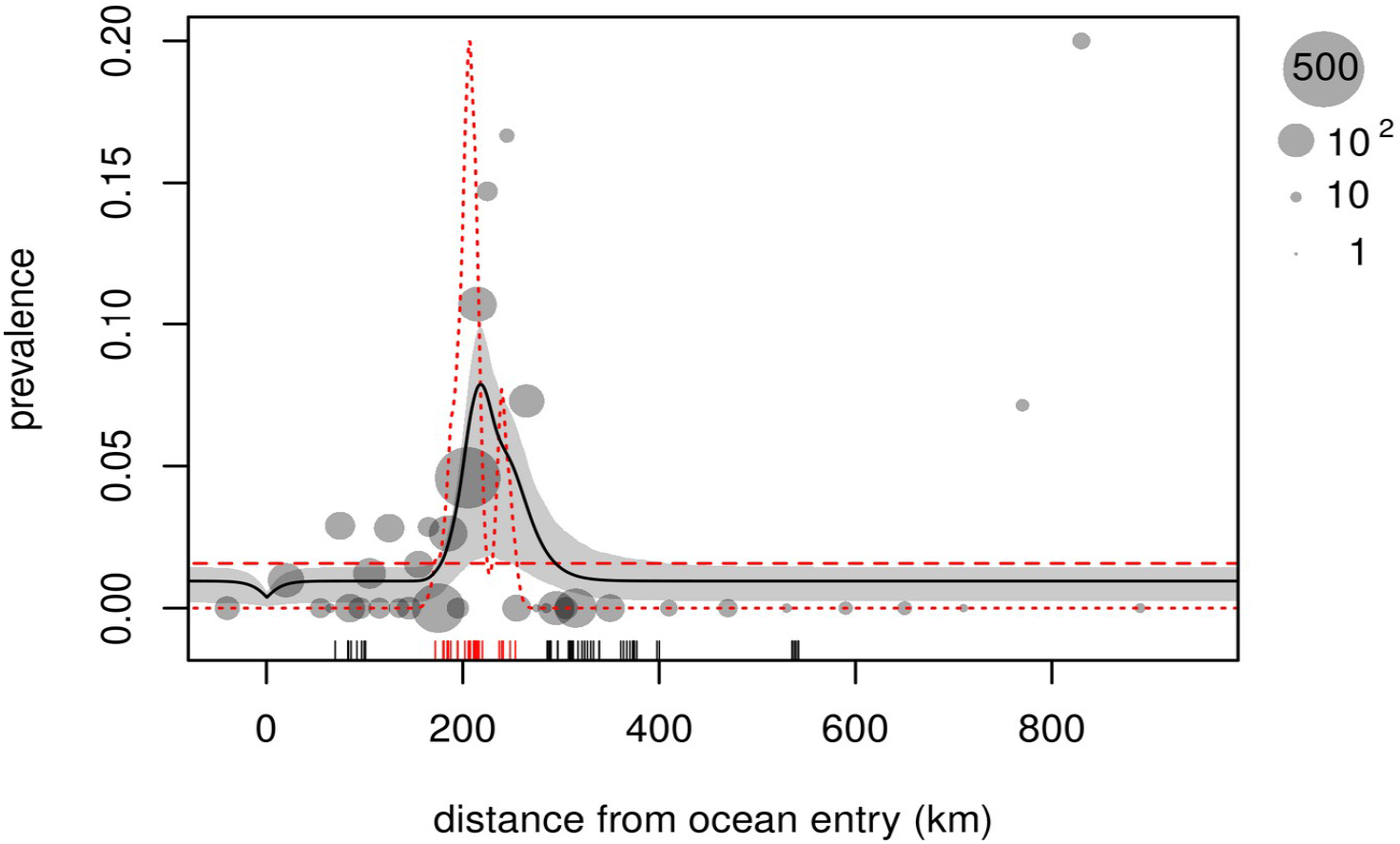
Prevalence of *Tenacibaculum maritimum* infection in Fraser-River sockeye salmon smolts migrating north-westward from the mouth of the Fraser River, BC (*x*=0). Grey circles represent smolts caught in trawl surveys and purse-seine sampling between 2008 and 2018 (aggregated by location to illustrate pattern). Black curve shows predictions from a spatio-epidemiological model of migration and infection dynamics, fitted to the prevalence data and plotted for the average Julian day of fish capture; surrounding light grey shows 95% confidence region, for model incorporating background and Discovery-Island-farm infection sources. Red curves show model-fit relative infection pressure from background (dashed) and salmon-farm-origin (dotted) sources, primarily in and around the Discovery Islands (scale arbitrary). “Rug” shows location of salmon farm tenures, with red indicating Discovery Islands farms.

The best model describes a scenario in which sockeye salmon smolts are subject to *T. maritimum* infection from both background and salmon-farm sources. Peak contributions from Discovery Island farms dwarfed those from other farming locations, and farm-source infection pressure peaked at 12.7 times background infection pressure. Despite this higher estimated peak exposure due to salmon farms, cumulative exposure along the migration route was higher for background sources. Numerically integrating background- and farm-source exposure rates across the migration window, farm-origin exposure occurred, over approximately a 100 km portion of the migration route, in approximately 0.21 of the modelled sockeye population. Background-origin exposure occurred in 0.43 of the sockeye population, over the entire 1600 km spatial window we modelled. (Note that some repeat exposure is possible due to recovery in the model.) The MLE parameters resulted in 67.7% survival (95% CI: 43.9% to 93.5%) by the end of the migration period, relative to survival that would result in the context of background infection only (Figure 5). That is, between 6.5% and 56.1% of sockeye smolts appear to die as a result of farm-origin *T. maritimum* exposure. (Note that confidence regions for survival in the full model and the background-only model [Figure 5] appear to overlap, but these rates are not independent; survival in the full model at most equals background-only survival.)

**Figure 5.**
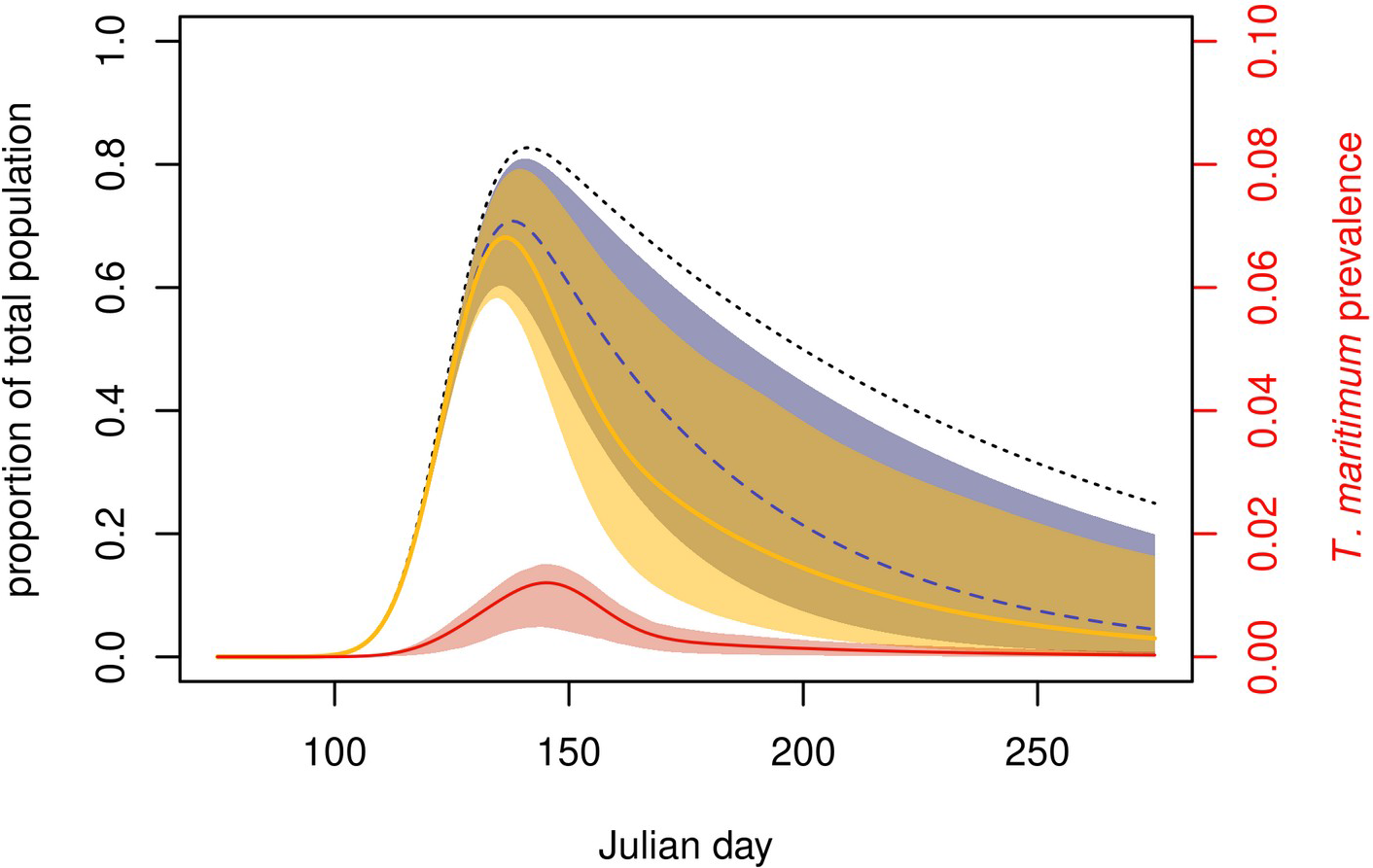
Spatio-epidemiological model predictions of proportional Fraser-River sockeye salmon smolt marine abundance (black) by Julian day, assuming 1: background-only mortality (dotted curve), 2: mortality due to background *Tenacibaculum maritimum* infection (dashed blue curve and 95% confidence region), and 3: mortality due to background and farm-origin *T. maritimum* infection (solid orange curve and 95% confidence region). Sockeye enter from freshwater over time but immediately incur some level of mortality, such that the whole population is never in saltwater at once and proportional abundance never reaches one. Red curve (and 95% confidence region) indicates *T. maritimum* infection prevalence in model 2 (note different scale).

**Figure 6.**
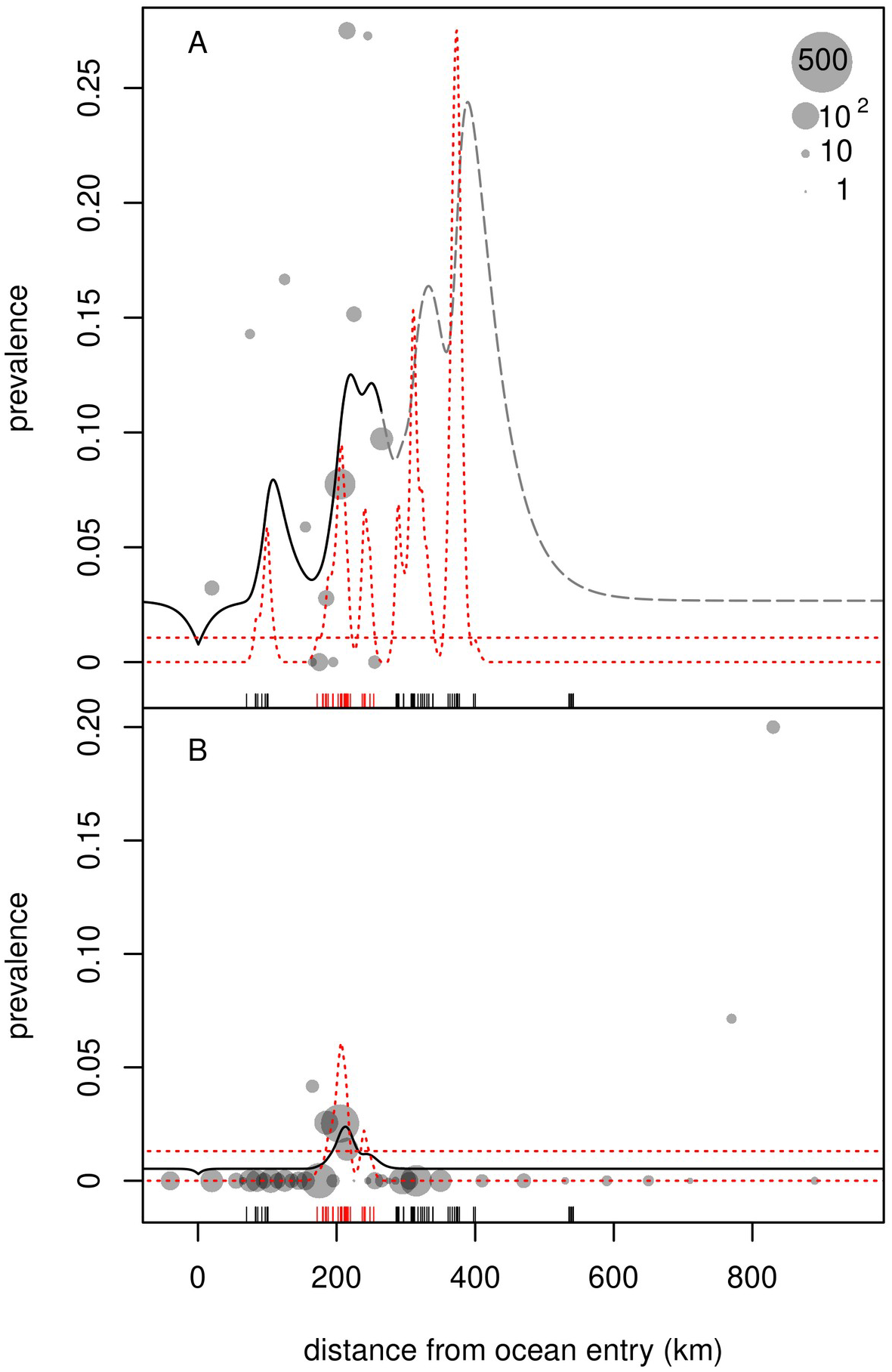
Prevalence of *Tenacibaculum maritimum* infection in Fraser-River sockeye smolts migrating northwest from the Fraser River, BC (*x*=0), for 2015 (A) and for all other study years from 2008 through 2018 (B). Grey circles represent smolts caught in trawl surveys and purse-seine sampling, aggregated by location to illustrate pattern. Black curves show predictions from spatio-epidemiological partial-differential-equation models of migration and infection dynamics, fitted to the relevant prevalence data and plotted for the average Julian day of fish capture. Model curve in dashed where data lacking. Red dashed lines show relative infection pressure from fitted constant and salmon-farm-origin sources in the corresponding time periods (scale arbitrary but matching in A and B). “Rug” shows location of salmon farm tenures, with red indicating Discovery Islands.

### Model uncertainty

Although there was substantial uncertainty in parameters from our best model, results indicated several clear features relevant for Fraser-River sockeye. The exponent for Discovery-Island-farm infection scaling factor, δ, was clearly greater than zero (Figure S2A) with a lower 95% confidence bound of 1.53, yielding a lower 95% confidence bound on the scaling factor itself, *e^δ^*, of 4.6. The best-fit value for *e^δ^* was 8.8×10^4^, however, and the likelihood appeared to decline monotonically with increasing δ. The standard deviation parameter for salmon-farm-origin infection pressure, *σ*, was estimated to be near the minimum value of 0.1 (Figures S2B, S4), with an upper 95% confidence bound of 12.8. A wide range of mortality (*μ_I_*) and recovery (*γ*) values produced similar model fits (Figure S2C), with the model unable to discern whether infected mortality was elevated, relative to baseline mortality, or whether infected individuals recovered at a non-zero rate.

Background and peak farm-origin and infection rates (*β*_0_ and *β_F_*, respectively) also displayed a wide range of likely values (Figure S2D), but the estimates indicate that farm-source infection pressure was 12.7 times background levels with a 95% confidence region that spanned 4.5 to 31 times background levels.

Even though infection, recovery, and associated mortality parameters were not simultaneously estimable, given the data, the resulting model showed a well resolved peak in *T. maritimum* prevalence near Discovery-Island salmon farms (Figure 4). The model form used to estimate uncertainty illustrated in Figure 4, however, considered only Discovery-Island salmon farms due to optimisation constraints. Figure S5 shows model predictions from the best-fit version of our most parsimonious model form (Model 8 from Table 2) with δ fixed at 1.53, the lower bound of its 95% confidence interval. Despite lingering uncertainty in the particulars of our best model, the influence of Discovery Islands farms stands out as receiving the overwhelming majority of model support across the models we considered (Table 2).

### Inter-annual variation

*T. maritimum* prevalence was high in 2015, and our best model, fit to data from that year alone, was consistent with much higher levels of background and farm-origin infection than the model fit to data from all other years (Figure 5A). The fitted model for 2015 indicates more even contributions from salmon farms along the sockeye migration route, due to higher rates of *T. maritimum* detection in fish caught further to the south. Unfortunately, sampling was spatially restricted in 2015, precluding validation of the putative pattern using samples from further north. Regardless, the infection peak associated with Discovery-Island farms did not disappear when 2015 data were omitted from those used to fit the model (Figure 5B).

### Information from salmon farms

Using our best model as a starting point, we considered two further model modifications, based on information from salmon farms.

First, *T. maritimum* in Atlantic salmon in BC tends to cause acute disease from which individuals can recover, not commonly displaying signs of disease again (DFO 2020). As a result, we fit an “SEIR” (susceptible exposed infected recovered) version of the best model (Equation S2), which allows for a new recovered class relative to the SIS and SEIS models. The likelihood was within 0.02 of the best-fitting model, with an equal number of parameters, presenting a nearly identical model fit (Figure S6). This result was consistent with our uncertainty calculations, which indicated that the modelling framework cannot resolve the details of recovery from infection (Figure S2C).

While farmed salmon do not tend to experience recurrent mouthrot, the initial bout on a farm can lead to substantial mortality, and fish are treated with antibiotics (Frisch et al. 2018). Treatment with appropriate antibiotics may therefore be an indicator of farm outbreaks during which *T. maritimum* release would be elevated (Z. Waddington and D. Price, personal communication 2020). We considered an additional modification to the best-fitting model, scaling treated farms contributions to farm-source infection pressure (Equation 2) by an additional free parameter in the months during which they were treated. Thus farms that were treated more often during the study period had the potential to contribute more to farm-source infection pressure. This model did not significantly improve the model fit (likelihood ratio test, p=0.36).

## Discussion

Fraser-River sockeye smolts, migrating northwest after ocean entry, displayed a sharp peak in *Tenacibaculum maritimum* detections in the Discovery Islands region of BC, about 200 km after they had left freshwater (Figure 2). Standard differential-equation models fit to these data indicate that Atlantic salmon farms in the Discovery Islands likely contribute to *T. maritimum* infection rates in Fraser-River sockeye. An advection-diffusion-decay sub-model, parameterised with data from multiple sources, described sockeye migration north from the Fraser River, and we fit competing SEIS (susceptible, exposed, infected, susceptible) sub-models to qPCR molecular-detection data to describe the epidemiology of *T. maritimum* in sockeye. Our best models, directly informed by the data, described a peak in farm-source infection that was an order of magnitude (12.7 times) higher than background infection rates. Resulting farm-origin infection occurred over a short portion of the early marine migration route, but at such high intensity that total farm-source infections approached total background-origin infections acquired over the entire migration route.

Using the data available, uncertainty in the model parameters left ambiguity around several features of interest, but the fitted models nevertheless produced reasonably precise spatiotemporal predictions. In particular, we could not make high-precision estimates for a number of parameters on their own. These uncertain parameters included: the weighting of infection pressure from Discovery-Island farms relative to other farms along the sockeye migration route, specific infection rates from farm and background sources, and recovery and mortality rates in infected fish (Figure S2). Due to the specific mathematical formulation of our model, the most we could say about Discovery-Island farm weighting is that it is high; farm-source *T. maritimum* infection pressure from Discovery-Island farms appears to far exceed that from other farms. In the case of farm- versus background-origin infection rates, the individual parameter values were uncertain but results indicate that peak farm-source infection pressure was between 4.5 and 31 times as high as average background infection pressure (Figure S2D). The question of whether infection prevalence declines due to mortality or recovery was particularly unclear, even though the maximum-likelihood parameter values describe a scenario in which most of the decline in prevalence was due to mortality. Regardless of uncertainties, the range of parameter values supported by the data all led to the same overall conclusions: at its peak, infection pressure from farm-origin sources appears to have been many times stronger than from background-origin sources, and the main peak of infection occurred near the cluster of farms in the Discovery Islands.

The fitted model indicates that *T. maritimum* infection can be detected shortly after farm-source exposure, resulting in peak infection close to salmon farms in the Discovery Islands (Figure 4). Although this progression from exposure to detectable infection is rapid, the timeline aligns with evidence from Atlantic salmon farms, which suggests that *T. maritimum* infection, disease development, and resulting mortality in untreated fish can be swift, with associated deaths observed as early as two days after transfer to saltwater (Frisch 2018). Even though evidence from another species cannot validate our sockeye model, per se, our model’s consistency with observations in another salmonid is reassuring. Known, rapid migration times for juvenile sockeye through the Discovery Islands (generally between two days and two weeks; Rechisky et al. 2021) further align with the description of directed migration in our model. Thus, an ability to rapidly detect *T. maritimum* in sockeye after they were exposed, paired with the smolts’ directional migration behaviour, appears to have facilitated identification of infection sources.

We note that the sockeye smolts we classified as infected may combine truly infected individuals with those merely exposed to *T. maritimum* at high loads. After such exposure, bacteria adhering to an individual’s gills, but not truly infecting the fish, might have been detected in our mixed-tissue screening procedure. Scepticism has been levelled at past uses of gill-inclusive mixed-tissue samples for screening on the Fluidigm BioMark™ platform (personal communication, various sources). Gills are, however, one of the tissues in which mouthrot manifests in Atlantic salmon (Frisch et al. 2018*b*), infections can become systemic (Frisch et al. 2018*a*), and our own assessments of single-tissue results (not shown) indicate that detections are indeed often systemic, especially when nucleic-acid copy numbers are high. We cannot exclude the possibility that the rapid increase in and subsequent decline of *T. maritimum* prevalence near Discovery-Island farms may be due to misclassification of exposure as infection, with “recovery” simply due to elution of bacteria into the marine environment. Regardless, Fraser River sockeye infected with or exposed to *T. maritimum* are most prevalent near salmon farms in the Discovery Islands, implying that those farms are – at a minimum – likely sources of exposure.

While we cannot attribute specific *T. maritimum-*induced mortality rates based on our analysis, elevated infection-induced mortality rates do align with other sources of information. In addition to the rapid mortality observed in farmed Atlantic salmon (Frisch 2018), mortality of lab-reared Atlantic salmon smolts cohabiting with artificially infected individuals has reached as high as 100% in ten days, for certain isolates of *T. maritimum* (Frisch et al. 2018*b*). Although a certain level of virulence in one host species does not directly imply similar virulence in another, it seems that a high rate of *T. maritimum-*induced mortality in affected sockeye would be reasonable to expect, especially considering that wild sockeye smolts are subject to additional foraging and predator-avoidance burdens, not present for farmed or lab-reared fish (Bakke and Harris 1998; Miller et al. 2014; DFO 2020), and that the region of peak infection imposes particular foraging constraints on salmon smolts (James et al. 2020). That said, our maximum-likelihood estimate of infection-induced mortality is likely too high, since resulting survival at the end of our roughly five-month migration window fell to 3.0% (95% CI: 0.2% to 16.4%), a level comparable to overall lifetime marine survival estimated for sockeye salmon (e.g. Irvine and Akenhead 2013).

Even without a definitive estimate of infection-induced mortality rates, our model results clearly illustrate the general point that infection prevalence need not be high to cause substantial mortality. The best-fit model parameters indicate a high mortality rate and relatively low recovery rate in infected individuals (Figure S2C), combining to reproduce the observed decline in infection prevalence as sockeye smolts migrated past the Discovery Islands (Figure 4). The resulting predicted mortality due to *T. maritimum was* 87.9% by Julian day 275 (150 days after mean ocean-entry date), even though population-wide infection prevalence peaked at just 1.2% around Julian day 150 (25 days after ocean entry). Although the fitted model indicates that many fish became infected, they either died as a result or recovered so that population-wide infection prevalence at any single point in time remained low. Apparent low-prevalence infections can belie high resultant mortality.

Our best fitting model’s counter-intuitive mismatch between low overall infection prevalence and high associated mortality is, in part, a result of external-source infection pressure, whereby the modelled sockeye population itself did not have to sustain and transmit the infection internally. On reflection, we should perhaps not expect high infection prevalence for a virulent pathogen in an impacted wild population if infections primarily derive from a stable external infection source. In general, direct disease-induced mortality can be rapid for some pathogens (e.g. Frisch et al. 2018*b*), and diseased individuals may suffer indirect mortality through their inability to find food or evade predators (Bakke and Harris 1998; Miller et al. 2014; James et al. 2020; Furey et al. 2021). As a result, infected individuals may drop out of a population without spending much time infected, keeping the overall proportion of infected individuals at any one time low. A key point here is that the pathogen does not have to sustain itself in the wild population, if a nearby stable infection source is present. Outbreaks that would normally “burn out” in the wild have the potential to persist in the system overall, and wild populations can be driven to extinction (de Castro and Bolker 2004). In the case of parasitic infections, this phenomenon has been described as an induced Allee effect that reservoir hosts can impose on endangered wildlife (Krkošek et al. 2013). There is no guarantee that any given rare pathogen will have a large effect, but in the scenario we describe a pathogen need not be common in a focal population to cause substantial losses. The data we analysed here are consistent with an effect in sockeye due to *T. maritimum* from Atlantic salmon farms that is disproportionate to the observed prevalence.

### Persistent Infection on salmon farms

Longitudinal data from salmon farms in BC indicate that Atlantic salmon can remain infected or become reinfected with *T. maritimum* throughout an 18-month production cycle (Bateman et al. 2021), but Atlantic salmon rarely, if ever, develop clinical mouthrot twice (Z. Waddington and D. Price, DFO Aquaculture Management veterinary staff, personal communication 2020). Not knowing how infection plays out in sockeye, we considered a “SEIR” version of our best-fitting model that included a “recovered” class of individuals that could not become reinfected. Results & model fit were practically identical to those of our most parsimonious model (Figure S6). This finding aligns with the conclusion that our modelling framework cannot distinguish between recovery and mortality that both serve to reduce infection prevalence. Such a result is common in epidemiological studies, and more detailed data on recovery versus mortality arising from direct and indirect ecological processes are needed.

While antibiotic treatment is almost certainly related to disease state in farmed fish, our lack of evidence for a treatment effect on farm infection pressure aligns with observations about *T. maritimum* from other studies (Laurin et al. 2019; Shea et al. 2020). In preliminary analyses (results not shown) we found no association between farm treatment status and either *T. maritimum* detection prevalence in fish sampled as part of DFO’s audit program or *T. maritimum* detections via eDNA sampling in the nearby marine environment. Further, although on-farm antibiotic treatment can extend for several months (Waddington and Price, personal communication 2020), dead and dying fish on farms can display elevated *T. maritimum* detection loads for somewhat longer, after any reported mouthrot outbreak has passed (Bateman et al. 2021). Together, these results suggest that antibiotic treatment, infection status, and possibly infectivity may not be intimately tied. Thus, farmed salmon infected with *T. maritimum* may continue to pose a risk to wild salmon even after the associated disease is brought under control on a farm.

### Environmental effects

Environmental features we did not consider likely contribute to some of the patterns we observed. We have not taken detailed ocean circulation patterns into account, although circulation models do not suggest that the Discovery Islands represent a region of natural pathogen retention (Foreman et al. 2015; Khangaonkar et al. 2017). The Discovery-Islands region does, however, act as a funnel for out-migrating juvenile sockeye as they leave the wider reaches of the Salish Sea to the south. This concentrates sockeye smolts in a confined space within which interactions with wild species may be amplified. *T. maritimum* is present in cnidarian species in other parts of the world (Delannoy et al. 2011), although the extent to which this occurs in BC is unknown. There has been past consideration of infection in Pacific herring (Marty et al. 2010), observed to school with juvenile sockeye at times (Johnson et al. 2019), but screening within our lab (Miller unpublished data) detected mostly low levels in just eleven out of 378 Pacific herring samples. The bacterium is certainly present in other wild Pacific salmon (Miller et al. unpublished results), which can also school with sockeye and may constitute an additional source of exposure.

Another plausible explanation for the patterns we observed is the spatial patterning of environmental stressors that render fish more susceptible to disease caused by *T. maritimum* (Santos et al. 2019). Water temperatures in eastern parts of the Discovery-Islands, and especially in the Salish Sea further south, tend to be warm relative to surrounding regions (Chandler et al. 2017). This may cause thermal stress in migrating smolts. Sockeye may also start to experience food stress as they pass through the Discovery Islands (James et al. 2020), which could in turn facilitate opportunistic *T. maritimum* infection. Rather than discount salmon farms in the Discovery Islands as sources of exposure, however, nearby sources of environmental stress may help to explain why those farms, and not farms from other regions, appear to contribute disproportionately to infection patterns.

The pattern of elevated *T. maritimum* infection pressure around the Discovery Islands broadly holds across years, but we saw substantial inter-annual variation. Our results indicate that *T. maritimum* in sockeye was particularly high in 2015, a year of anomalously high ocean temperatures (Kintisch 2015). High infection rates in 2015, which served to drive smolt mortality predictions upwards in our overall model fit, correspond to particularly poor marine survival for Fraser sockeye that returned to spawn in 2017 (Hawkshaw et al. 2020). Ocean temperatures themselves are inversely related to sockeye marine survival, however (Hawkshaw et al. 2020), and *T. maritimum*, sometimes described as an “opportunistic” pathogen (Avendaño-Herrera et al. 2006), may be an indicator of other determinants of poor survival in smolts. Still, we might expect to see more *T. maritimum*-related issues for salmon farms, or more broadly, as ocean temperatures warm over the coming decades. On the other hand, high *T. maritimum* prevalence was not restricted to 2015; in 2012 a cluster of *T. maritimum* detections occurred around 800 km from the mouth of the Fraser River, and was not captured by any of the models we considered. Further investigation of inter-annual and spatial patterns seems warranted.

### Conservation implications

Overall, it is clear that detections of *T. maritimum* in Fraser River sockeye smolts are substantially elevated surrounding an intensive salmon-farming region of BC, the Discovery Islands. Salmon farms in this area have been the subject of intense scrutiny for over a decade, largely because of the potential disease risks they pose to sockeye and other wild salmon (Connors et al. 2012; Morton and Routledge 2016). The models and data we present are consistent with combined farm- and background-source *T*. *maritimum* infection and associated mortality in Fraser River sockeye. Salmon farms have long been tied, globally, to sympatric declines in wild salmonids (Ford and Myers 2008), and our analysis provides an in-depth case study that builds on previous high-level correlative investigation of Fraser River sockeye salmon survival in relation to salmon farm production in the Discovery Islands (Connors et al. 2012).

Scientific advice provided to Canadian policymakers identified *T. maritimum* as posing minimal risk to Fraser-River sockeye (DFO 2020). That assessment, however, did not access many of the data we used here, which indicate a spatial association between *T. maritimum* infection of juvenile sockeye salmon and salmon farms in the Discovery Islands. Taken together, results from our wild-salmon screening program highlight *T. maritimum* transmission as a possible mechanism for population-level impacts of farmed Atlantic salmon on wild Pacific salmon populations in BC, and present evidence that elevated infections in Fraser-River sockeye likely originate from salmon-farm sources in the Discovery Islands region. These findings, paired with the continued declines in sockeye population health (COSEWIC 2017; Hawkshaw et al. 2020), suggest that recent conclusions about the “minimal risk” of harm posed by this agent (DFO 2020) were premature.

During final preparation of this study, the Government of Canada announced a decision to remove open-net salmon farms from the Discovery Islands region of BC (https://www.canada.ca/en/fisheries-oceans/news/2020/12/government-of-canada-moves-to-phase-out-salmon-farming-licences-in-discovery-islands-following-consultations-with-first-nations.html). While this decision was made on the basis of consultations with local Indigenous title holders, rather than risks posed by salmon aquaculture per se, our findings suggest that the decision was nonetheless precautionary in effect, when viewed from the perspective of conserving Fraser-River sockeye salmon.

### Broader conclusions

Sockeye salmon are just one example of the many migratory species that globally travel large distances to complete their life cycles. Humanity’s footprint on planet Earth is ever expanding (Ripple et al. 2017), and the wildlife-livestock interface will likely continue to gain relevance for a growing proportion of migratory species, as components of their habitat face agricultural incursions. Within this context, negative interactions between wild species and nearby livestock are likely to increase (Gordon 2018), with disease transfer risks providing just one example. Further, the unprecedented rate of species domestication for aquaculture, in particular (Duarte et al. 2007), and a suite of other risk factors for disease transfer in the marine environment (Krkošek 2017), mean that the marine wildlife-livestock interface will likely remain a hotspot for infectious disease in migratory marine species and a focal point for future research.

Migratory species are particularly difficult to conserve, given that they rely on multiple connected but geographically distinct habitat regions (Runge et al. 2014). Despite optimistic examples of conserving migratory corridors (Purdon et al. 2018; Chester and Hilty 2019), maintaining adequate connected habitat remains a challenge, even in the wealthiest countries (Berger et al. 2014). While not a physical barrier to movement, infectious diseases from livestock could pose similar challenges for obligate migratory species, reducing the proportion of a population able to successfully migrate. Species would have to adapt (e.g. immunologically, phenologically, or behaviourally) or perish. For ocean-run sockeye salmon, to use our focal example, adaptation could be possible, since the species has repeatedly evolved a landlocked form (kokanee; Wood et al. 2008) and a small proportion of Fraser-River sockeye already use a less salmon-farm-exposed southern migration route (Tucker et al. 2009; Morton and Routledge 2016). Many other species might not be so lucky (Hardesty-Moore et al. 2018). For those species that evolved migration strategies in response to disease pressure, migration barriers – disease-induced or otherwise imposed – could even trap populations in the same maladaptive scenarios they originally evolved to evade (Satterfield et al. 2015).

If migration – clearly an adaptive strategy prior to human interventions – is to persist, we must ensure the necessary integrity of all components of migratory species’ requisite habitats. Much greater precaution and coordinated action will almost certainly be necessary (Mason et al. 2020).

## Acknowledgements

We are very grateful to the Pacific Salmon Foundation and Genome British Columbia for funding and support to carry out the Strategic Salmon Health Initiative, the overarching program under which this study operated. This program was further part of the Salish Sea Marine Survival Project, led by the Pacific Salmon Foundation under the scientific leadership of Dr. Brian Riddell and Dr. Isobel Pearsall. MK is grateful for a NSERC Discovery Grant, Accelerator Award, and Canada Research Chair. Thanks to Marc Trudel, Chrys Neville, Jackie King, and the Hakai Juvenile Salmon Program for providing samples. Thanks to Mark Lewis, Marie Auger-Méthé, Brendan Connors, and Cameron Freshwater for discussions at early stages of this analysis.

## Supplementary Information

**Table S1.** sockeye stocks analysed

| sockeye stocks | migration timing | citation |
| --- | --- | --- |
| Early Stuart, Rossette, Blackwater, Dust, Paula, Cayenne | Early Stuart | (Neville et al. 2016) |
| Bowron, Chilliwack, Dolly Varden, Fennel, Gates, Nadina, Nahatlatch, Pitt, Seymour, Scotch, Upper Adams | Early Summer | (Neville et al. 2016) |
| Chilko, Kuzkwa Creek, Pinchi Creek, Middle River, North Thompson, Quesnel, Raft, Stellako, Tachie | Summer | (Neville et al. 2016; COSEWIC 2017) |
| Cultus, Big Silver, Birkenhead, Late Shuswap, Lower Shuswap, Little Shuswap, Weaver | Late | (Neville et al. 2016; COSEWIC 2017) |

### Distance-to-farms as a function of migration distance

**Figure S1.**
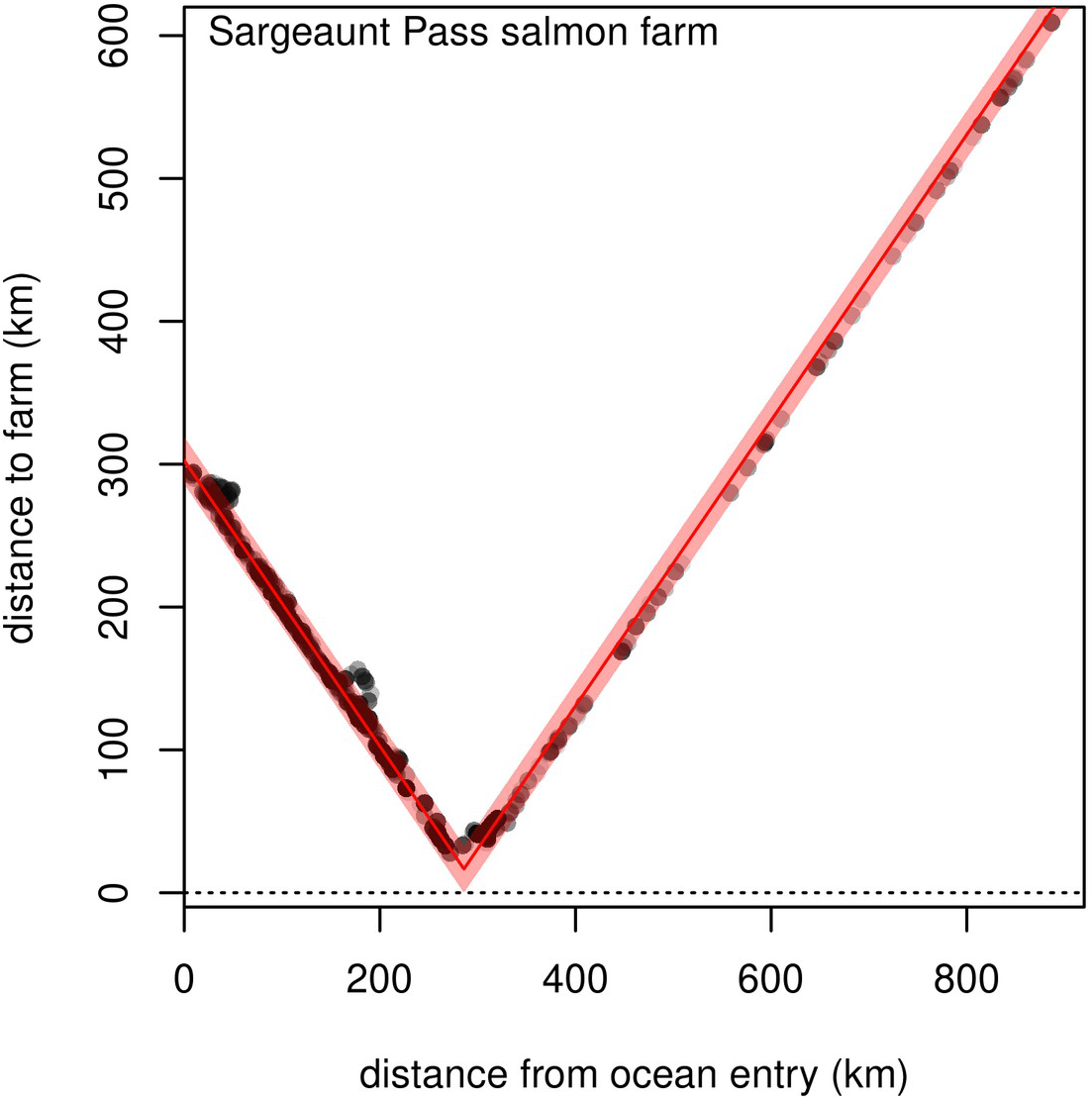
Example of relationship between migration distance and seaway distance to the Sargeaunt Pass salmon farm, for sampled sockeye salmon. The red line segments represent a fitted absolute-value model, with ordinate and abscissa shifts, and the shaded region represents 95% of the probability density associated with the normal distribution of model residuals.

### Additional model forms

The advection-diffusion-decay susceptible-only model (used for tuning the spatial component of the model) took the form:

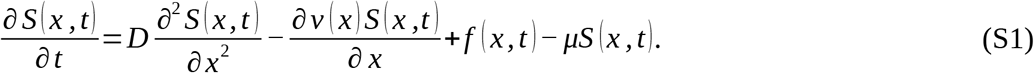

The SEIR version of the model described in the main text takes the form:

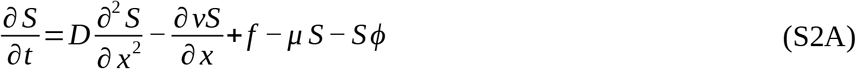

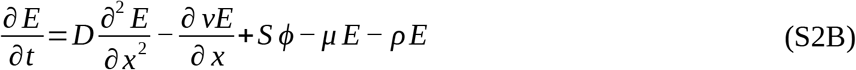

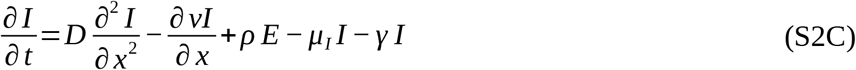

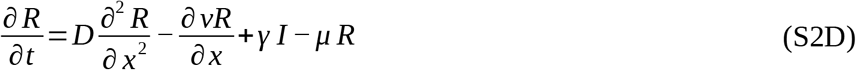

### Uncertainty

A likelihood profile for a single parameter can be generated by fixing the parameter of interest at a sequence of relevant values and, for each, minimising the model’s “restricted” negative log likelihood across the remaining free parameters. Twice the difference between the minimum value of the restricted negative log likelihood and the overall minimum negative log likelihood is asymptotically X^2^-distributed with one degree of freedom, allowing a confidence interval to be generated from the likelihood profile (e.g. Bolker 2008). The same process can be used to generate multidimensional likelihood profiles, and associated confidence regions, with X^2^ degrees of freedom equal to the number of restricted parameters.

**Figure S2.**
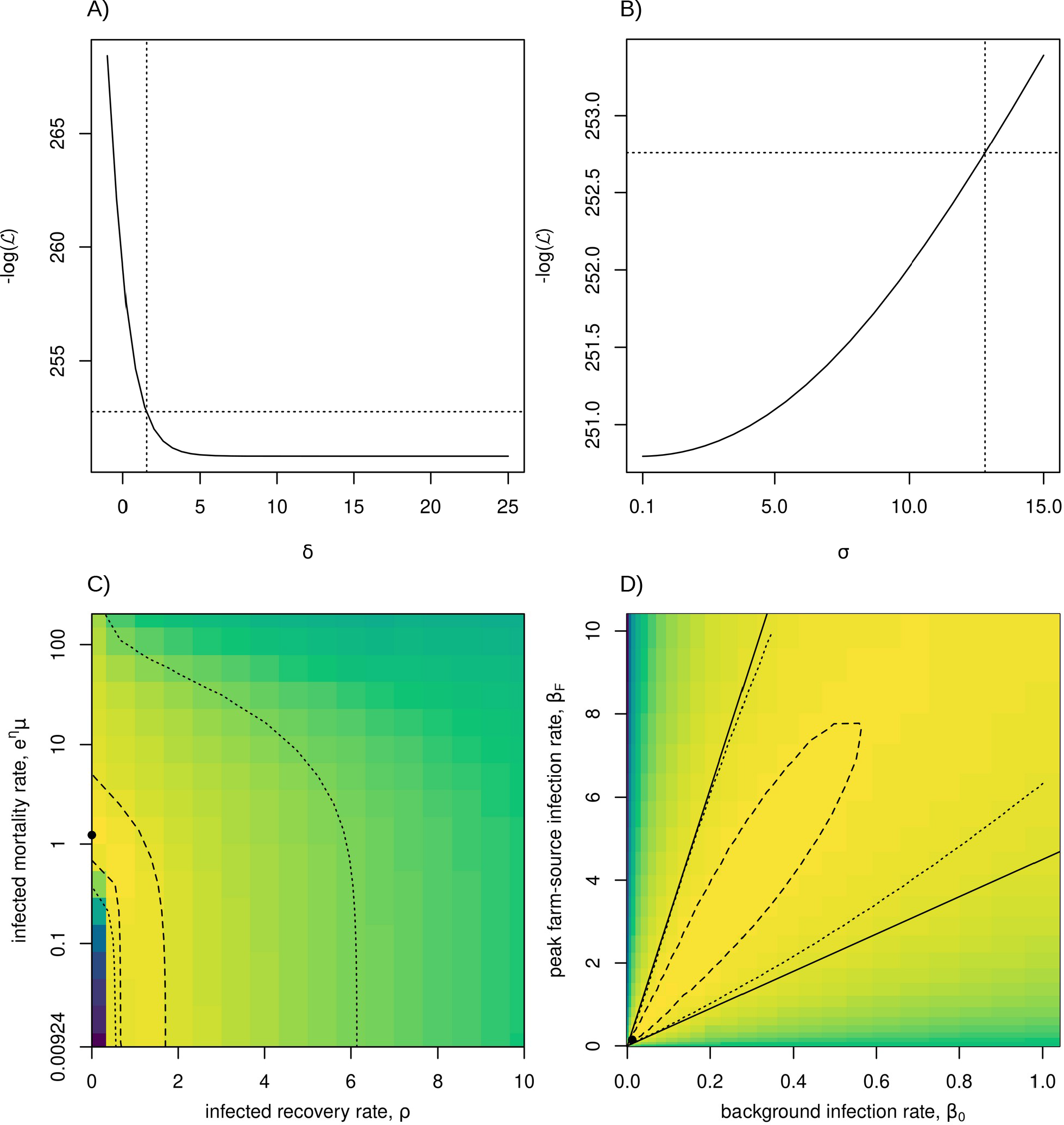
Likelihood profiles/surfaces and confidence intervals/regions for key model parameters. A) Likelihood profile for Discovery-Island infection weighting exponent (*δ*; solid curve) with dotted lines indicating 95% confidence interval (vertical) and associated critical likelihood value (horizontal). B) Likelihood profile for standard deviation of farm-infection kernel (*σ*) with 95% confidence interval and critical likelihood value indicated as in A). C) Two-dimensional likelihood surface infected mortality rate (*μ_I_* = *e^η^μ*; 0.00924 is baseline survival, *μ*) and infected recovery rate (*γ*). Yellow indicates lower values and contours indicates 95% (dotted) and 50% (dashed) confidence regions. Point shows maximum-likelihood parameter values (note log scale on mortality axis). D) Likelihood surface for background infection rate (*β*_0_) and maximum farm-source infection rate (*β_F_*) with confidence regions and maximum-likelihood values indicated as in C); solid lines have slopes of 4.5 and 31.

**Figure S3.**
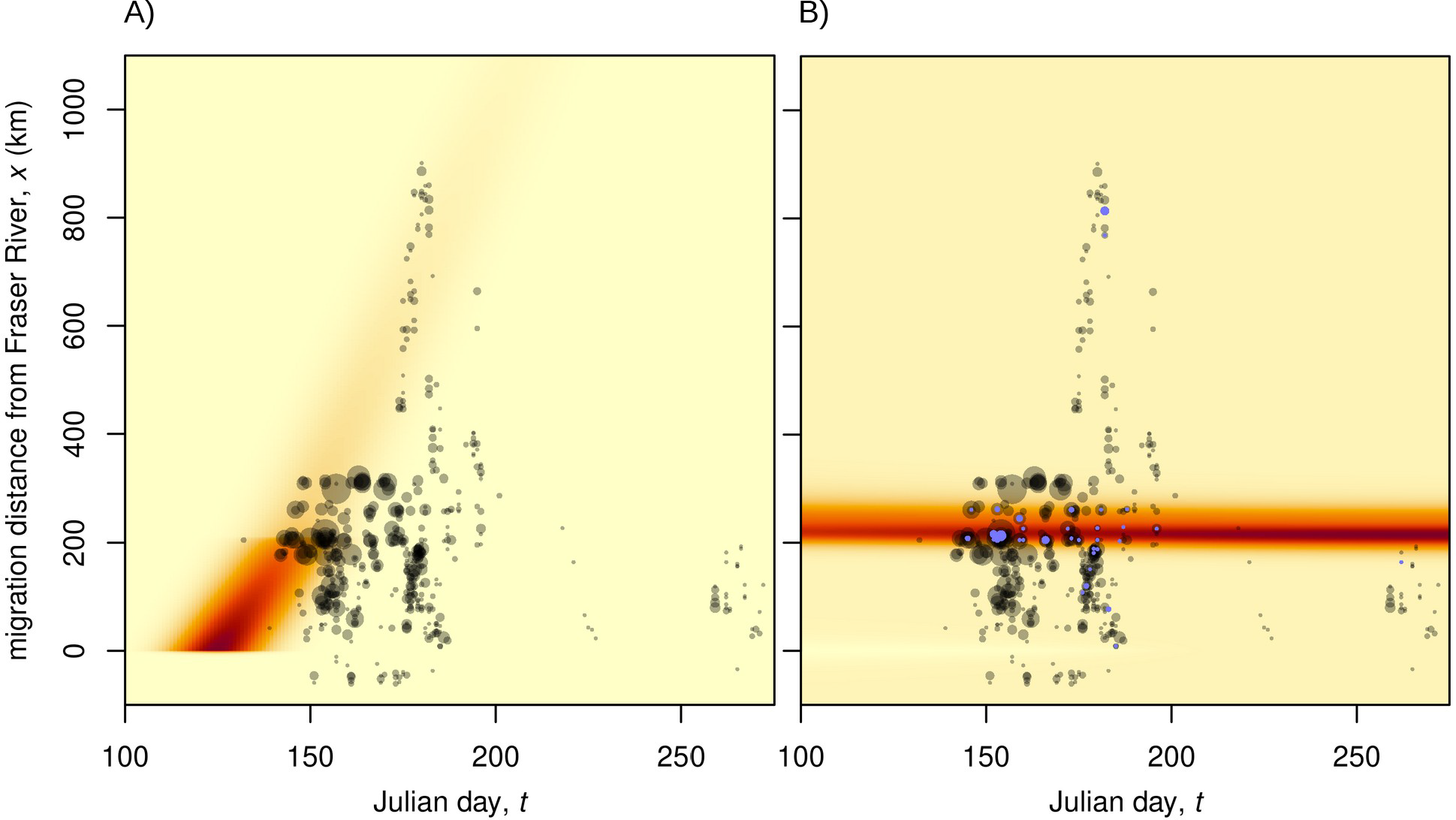
Model-output population density (A) and *Tenacibaculum maritimum* infection prevalence (B) in space and time. Darker regions indicate higher density or prevalence. Grey circles (area proportional to sample size) show sampling dates and locations for out-migrating Fraser-River sockeye, and corresponding blue circles in B) indicate sampled fish that tested positive for *T. maritimum*.

**Figure S4.**
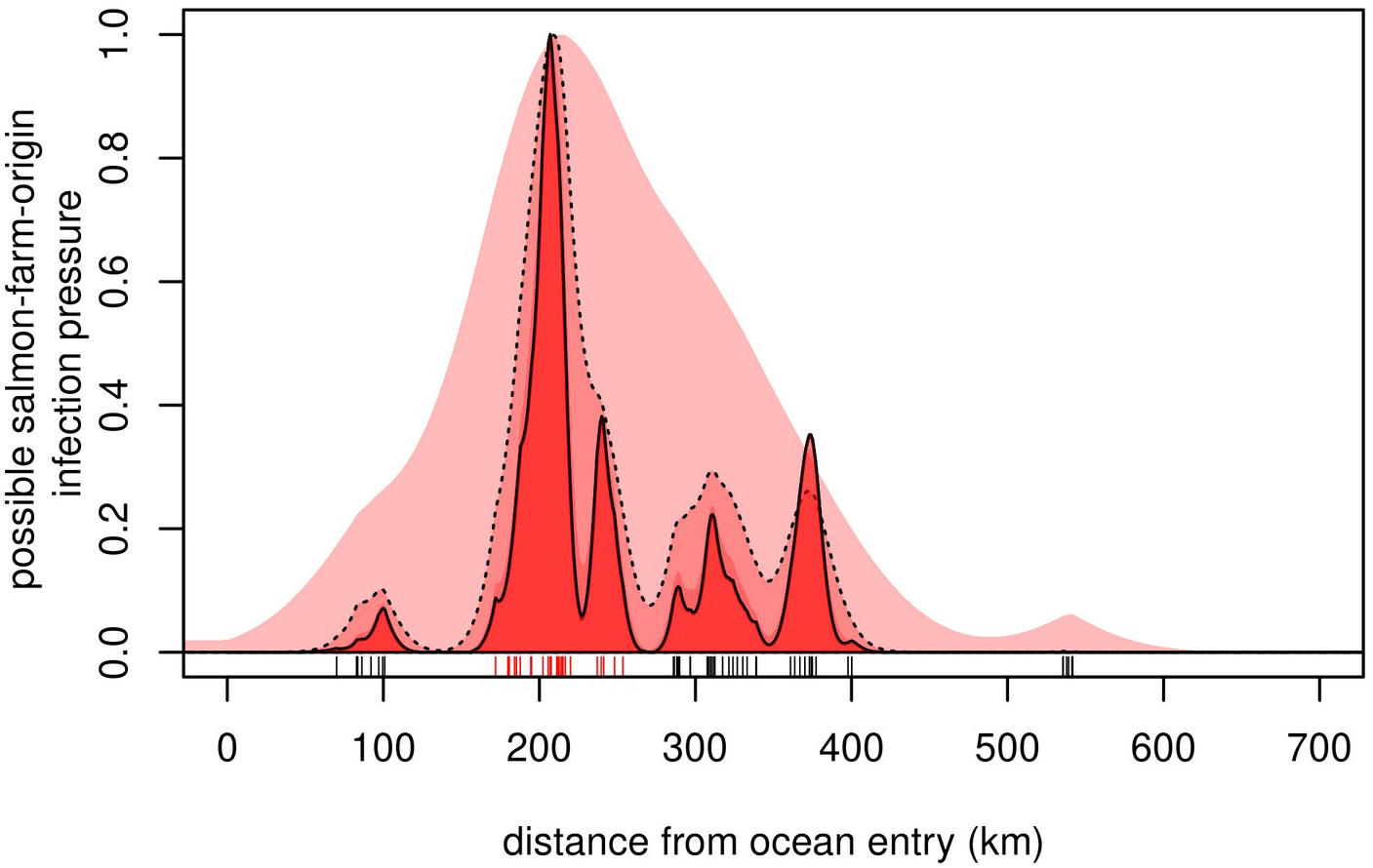
Combined salmon-farm-origin *Tenacibaculum maritimum* infection pressure in northwest-migrating Fraser River sockeye smolts for different values of the “spread” parameter, *σ* (standard deviation of single-farm Gaussian dispersal kernel). From lightest to darkest, plot shows *σ* equal to 50, 12.8 (dashed curve; 95% CI bound for the most parsimonious model in the main text), 5, 1 (solid curve, MLE value from the most parsimonious model), and 0.1. For illustration purposes, contributions from all farms are scaled by farm activity, and do not vary by region. Estimates resulting from *σ* values of 1 and 0.1 almost completely overlap and are indistinguishable in the plot. “Rug” shows location of salmon farm tenures, with red indicating Discovery Islands.

**Figure S5.**
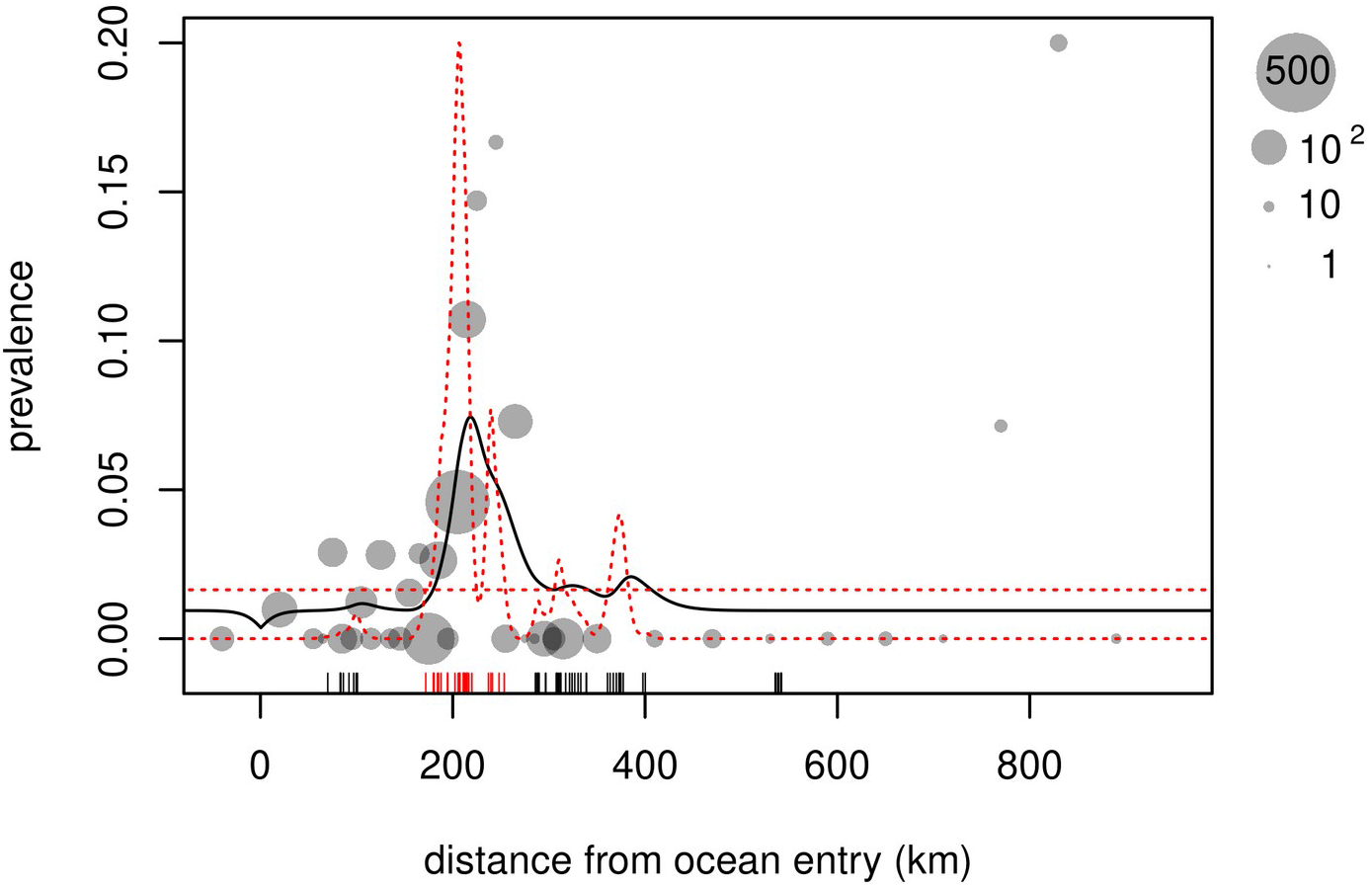
Prevalence of *Tenacibaculum maritimum* infection in Fraser-River sockeye salmon smolts migrating north-westward from the mouth of the Fraser River, BC (*x*=0). Grey circles represent smolts caught in trawl surveys and purse-seine sampling between 2008 and 2018 (aggregated by location to illustrate pattern). Black curve shows predictions from a spatio-epidemiological model of migration and infection dynamics, fitted to the prevalence data and plotted for the average Julian day of fish capture. Red dashed lines show model-fit relative infection pressure from background and salmon-farm-origin sources (scale arbitrary); weighting of infection pressure from Discovery-Island farms, relative to farms from other regions, is fixed at 4.60 (the lower bound of the associated 95% confidence interval). “Rug” shows location of salmon farm tenures, with red indicating Discovery Islands farms.

**Figure S6.**
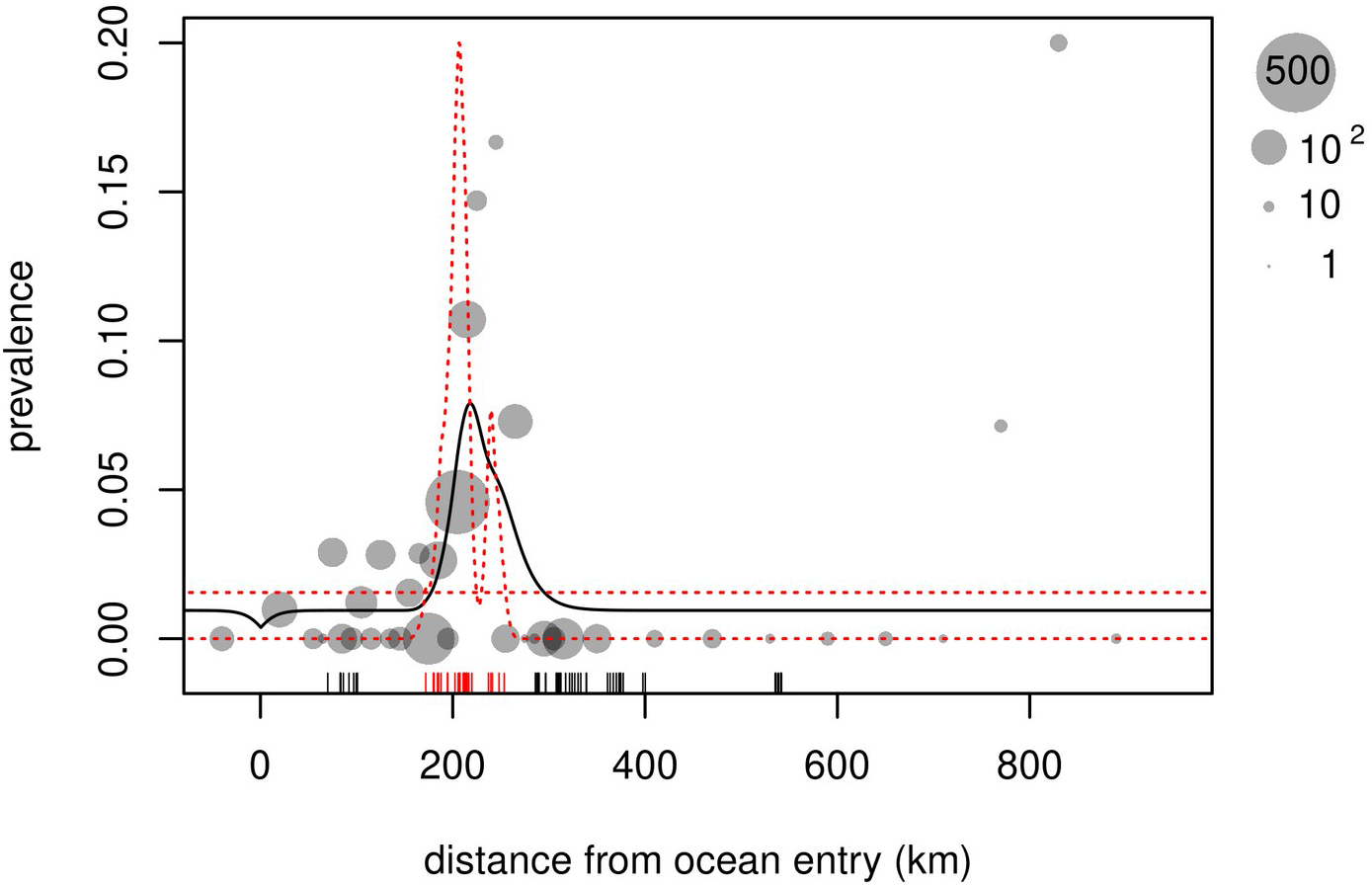
Prevalence of *Tenacibaculum maritimum* infection in Fraser-River sockeye salmon smolts migrating north-westward from the mouth of the Fraser River, BC (*x*=0). Grey circles represent smolts caught in trawl surveys and purse-seine sampling between 2008 and 2018 (aggregated by location to illustrate pattern). Black curve shows predictions from a spatio-epidemiological model of migration and infection dynamics, allowing for full recovery from infection, fitted to the prevalence data and plotted for the average Julian day of fish capture. Red dashed lines show model-fit relative infection pressure from background and salmon-farm-origin sources (scale arbitrary). “Rug” shows location of salmon farm tenures, with red indicating Discovery Islands farms.

## Notes

### Competing Interest Statement

The authors have declared no competing interest.

## References

Akaike, H. 1973. Information theory and an extension of the maximum likelihood principle. Pages 268–281 *in* B. N. Petrov and F. Csaki, eds. Second International Symposium on Information Theory. Akadémiai Kiadó, Budapest.

Altizer, S., R. Bartel, and B. A. Han. 2011. Animal migration and infectious disease risk. science 331:296–302.

Avendaño-Herrera, R., A. Toranzo, and B. Magariños. 2006. Tenacibaculosis infection in marine fish caused by Tenacibaculum maritimum: a review. Diseases of Aquatic Organisms 71:255–266.

Bakke, T. A., and P. D. Harris. 1998. Diseases and parasites in wild Atlantic salmon (Salmo salar) populations 55:20.

Bass, A. L., C. F. Stevenson, A. D. Porter, E. L. Rechisky, N. B. Furey, S. J. Healy, A. M. Kanigan, et al. 2020. In situ experimental evaluation of tag burden and gill biopsy reveals survival impacts on migrating juvenile sockeye salmon. Canadian Journal of Fisheries and Aquatic Sciences 77:1865–1869.

Bateman A. W., A. D. Schulze, K. H. Kaukinen, A. Tabata, G. Mordecai, K. Flynn, A. Bass, et al. 2021. Descriptive multi-agent epidemiology via molecular screening on Atlantic salmon farms in the northeast Pacific Ocean. Scientific Reports 11:3466.

Beacham T. D., M. Lapointe, J. R. Candy, B. McIntosh, C. MacConnachie, A. Tabata, K. Kaukinen, et al. 2004. Stock Identification of Fraser River Sockeye Salmon Using Microsatellites and Major Histocompatibility Complex Variation. Transactions of the American Fisheries Society 133:1117–1137.

Beacham T. D., B. McIntosh, and C. Wallace. 2010. A comparison of stock and individual identification for sockeye salmon (Oncorhynchus nerka) in British Columbia provided by microsatellites and single nucleotide polymorphisms. Canadian Journal of Fisheries and Aquatic Sciences 67:1274–1290.

Bengis R. G., R. A. Kock, and J. R. Fisher. 2002. Infectious animal diseases: the wildlife/livestock interface. Revue Scientifique et Technique de l’OIE 21:53–65.

Berger, J., S. L. Cain, E. Cheng, P. Dratch, K. Ellison, J. Francis, H. C. Frost, et al. 2014. Optimism and Challenge for Science-Based Conservation of Migratory Species in and out of U.S. National Parks: Conservation Action for Migratory Species. Conservation Biology 28:4–12.

Bolker, B. M. 2008. Ecological models and data in R. Princeton University Press, Princeton, NJ.

Burnham K. P., and D. R. Anderson. 2002. Model selection and multimodel inference: a practical information-theoretic approach. Springer Verlag, New York.

Chandler, P., M. G. G. Foreman, M. Ouellet, C. Mimeault, and J. Wade. 2017. Oceanographic and environmental conditions in the Discovery Islands, British Columbia.

Chester C. C., and J. A. Hilty. 2019. The Yellowstone to Yukon Conservation Initiative as an adaptive response to climate change. Pages 179–193 *in* Handbook of Climate Change and Biodiversity. Springer.

Cohen, B. I. 2012. The uncertain future of Fraser River sockeye. Commission of Inquiry into the Decline of Sockeye Salmon in the Fraser River (Canada). Commission of Inquiry into the Decline of Sockeye Salmon in the Fraser River, Vancouver, B.C.

Connors, B., and D. Braun. 2012. Migration links ocean-scale competition and local ocean conditions with exposure to farmed salmon to shape wild salmon dynamics. Conservation Letters.

Connors, B., D. Braun, R. Peterman, A. Cooper, J. Reynolds, L. Dill, G. Ruggerone, et al. 2012. Migration links ocean-scale competition and local ocean conditions with exposure to farmed salmon to shape wild salmon dynamics. Conservation Letters 5:304–312.

COSEWIC. 2017. COSEWIC assessment and status report on the sockeye salmon, Oncorhynchus nerka, 24 designatable units in the Fraser River drainage basin, in Canada.

Costello, M. J. 2009. The global economic cost of sea lice to the salmonid farming industry. Journal of Fish Diseases 32:115–118.

Csardi, G., and T. Nepusz. 2006. The igraph software package for complex network research. InterJournal, complex systems 1695:1–9.

de Castro, F., and B. Bolker. 2004. Mechanisms of disease-induced extinction. Ecology Letters 8:117–126.

Delannoy, C. M. J., J. D. R. Houghton, N. E. C. Fleming, and H. W. Ferguson. 2011. Mauve Stingers (Pelagia noctiluca) as carriers of the bacterial fish pathogen Tenacibaculum maritimum. Aquaculture 311:255–257.

DFO. 2020. Advice from the assessment of the risk to Fraser River Sockeye Salmon due to Tenacibaculum maritimum transfer from Atlantic Salmon farms in the Discovery Islands area, British Columbia (Science Advisory Report). DFO Canadian Science Advice Secretariat. National Capital Region.

Dijkstra, E. W. 1959. Dijkstra 1959 (shortest graph path).pdf. Numerische Mathematik 1:269–271.

Duarte C. M., N. Marbá, and M. Holmer. 2007. Rapid domestication of marine species. Science 316:382.

Ford J. S., and R. A. Myers. 2008. A global assessment of salmon aquaculture impacts on wild salmonids. PLoS biology 6:e33.

Foreman, M. G. G., M. Guo, K. A. Garver, D. Stucchi, P. Chandler, D. Wan, J. Morrison, et al. 2015. Modelling Infectious Hematopoietic Necrosis Virus Dispersion from Marine Salmon Farms in the Discovery Islands, British Columbia, Canada. PloS one 10:e0130951.

Frazer, L., A. Morton, and M. Krkošek. 2012. Critical thresholds in sea lice epidemics: evidence, sensitivity and subcritical estimation. Proceedings of the Royal Society B 279:1950–1958.

Fringuelli, E., P. D. Savage, A. Gordon, E. J. Baxter, H. D. Rodger, and D. A. Graham. 2012. Development of a quantitative real-time PCR for the detection of Tenacibaculum maritimum and its application to field samples: Real-time PCR for Tenacibaculum maritimum. Journal of Fish Diseases 35:579–590.

Frisch, K. M. 2018. Mouthrot in farmed Atlantic salmon.

Frisch, K., S. B. Småge, R. Johansen, H. Duesund, Ø. J. Brevik, and A. Nylund. 2018a. Pathology of experimentally induced mouthrot caused by Tenacibaculum maritimum in Atlantic salmon smolts. (P. Boudinot, ed.)PLOS ONE 13:e0206951.

Frisch, K., S. B. Småge, C. Vallestad, H. Duesund, ø J. Brevik, A. Klevan, R. H. Olsen, et al. 2018b. Experimental induction of mouthrot in Atlantic salmon smolts using *Tenacibaculum maritimum* from Western Canada. Journal of Fish Diseases 41:1247–1258.

Fritzsche McKay, A., and B. J. Hoye. 2016. Are Migratory Animals Superspreaders of Infection?: An Introduction to the Symposium. Integrative and Comparative Biology 56:260–267.

Furey N. B., A. L. Bass, K. M. Miller, S. Li, A. G. Lotto, S. J. Healy, S. M. Drenner, et al. 2021. Infected juvenile salmon can experience increased predation during freshwater migration. Royal Society Open Science 8:rsos.201522, 201522.

Gordon, I. J. 2018. Review: Livestock production increasingly influences wildlife across the globe. Animal 12:s372–s382.

Groot, C., L. Margolis (eds.)., and L. Margolis. 1991. Pacific Salmon Life Histories. UBC Press, Vancouver, B.C. (Vol. 564 pp.). UBC press, Vancouver, Canada.

Hardesty-Moore, M., S. Deinet, R. Freeman, G. C. Titcomb, E. M. Dillon, K. Stears, M. Klope, et al. 2018. Migration in the Anthropocene: how collective navigation, environmental system and taxonomy shape the vulnerability of migratory species. Philosophical Transactions of the Royal Society B: Biological Sciences 373:20170017.

Hastein, T., and T. Lindstad. 1991. Diseases in wild and cultured salmon: possible interaction. Aquaculture 98:277–288.

Hawkshaw M. A., Y. Xu, and B. Davis. 2020. Pre-season run size forecasts for Fraser River sockeye(Oncorhynchus nerka) and pink(O. gorbuscha) salmon in 2019. DFO, Ottawa, ON(Canada).

Irvine J. R., and S. A. Akenhead. 2013. Understanding smolt survival trends in sockeye salmon. Marine and Coastal Fisheries 5:303–328.

James S. E., E. A. Pakhomov, N. Mahara, and B. P. V. Hunt. 2020. Running the trophic gauntlet: Empirical support for reduced foraging success in juvenile salmon in tidally mixed coastal waters. Fisheries Oceanography 29:290–295.

Johnson, B., J. C. L. Gan, S. C. Godwin, M. Krkosek, and B. P. V. Hunt. 2019. Juvenile Salmon Migration Observations in the Discovery Islands and Johnstone Strait in British Columbia, Canada in 2018. North Pacific Anadromous Fish Commission 22.

Khangaonkar, T., W. Long, and W. Xu. 2017. Assessment of circulation and inter-basin transport in the Salish Sea including Johnstone Strait and Discovery Islands pathways. Ocean Modelling 109:11–32.

Kintisch, E. 2015. ‘The Blob’ invades Pacific, flummoxing climate experts. Science 348:17–18.

Krkosek, M. 2010. Host density thresholds and disease control for fisheries and aquaculture. Aquaculture Environment Interactions 1:21–32.

Krkošek, M. 2017. Population biology of infectious diseases shared by wild and farmed fish. Canadian Journal of Fisheries and Aquatic Sciences 74:620–628.

Krkošek, M., J. Ashander, L. N. Frazer, and M. a Lewis. 2013. Allee effect from parasite spill-back. The American naturalist 182:640–52.

Krkošek, M., B. M. Connors, A. Morton, M. A. Lewis, L. M. Dill, and R. Hilborn. 2011. Effects of parasites from salmon farms on productivity of wild salmon. Proceedings of the National Academy of Sciences 108:14700–14704.

Krkošek, M., M. A. Lewis, and J. P. Volpe. 2005. Transmission dynamics of parasitic sea lice from farm to wild salmon. Proceedings of the Royal Society B: Biological Sciences 272:689–696.

Laurin, E., D. Jaramillo, R. Vanderstichel, H. Ferguson, K. H. Kaukinen, A. D. Schulze, I. R. Keith, et al. 2019. Histopathological and novel high-throughput molecular monitoring data from farmed salmon (Salmo salar and Oncorhynchus spp.) in British Columbia, Canada, from 2011–2013. Aquaculture 499:220–234.

Maekawa, K., and S. Nakano. 2002. To sea or not to sea: a brief review on salmon migration evolution. Fisheries science 68:27–32.

Marty G. D., S. M. Saksida, and T. J. Quinn. 2010. Relationship of farm salmon, sea lice, and wild salmon populations. Proceedings of the National Academy of Sciences of the United States of America 107:22599–22604.

Mason, N., M. Ward, J. E. Watson, O. Venter, and R. K. Runting. 2020. Global opportunities and challenges for transboundary conservation. Nature ecology & evolution 4:694–701.

Miller K. M., I. A. Gardner, R. Vanderstichel, T. Burnley, A. D. Schulze, S. Li, A. Tabata, et al. 2016. Report on the Performance Evaluation of the Fluidigm BioMark Platform for High Throughput Microbe Monitoring in Salmon. Fisheries and Oceans Canada Canadian Science Advisory Secretariat.

Miller K. M., S. Li, K. H. Kaukinen, N. Ginther, E. Hammill, J. M. Curtis, D. A. Patterson, et al. 2011. Genomic signatures predict migration and spawning failure in wild Canadian salmon. science 331:214–217.

Miller K. M., A. Teffer, S. Tucker, S. Li, A. D. Schulze, M. Trudel, F. Juanes, et al. 2014. Infectious disease, shifting climates, and opportunistic salmon declines. Evolutionary Applications.

Morton, A., and R. Routledge. 2016. Risk and precaution: Salmon farming. Marine Policy 74:205–212.

Murray, J. D. 2007. Mathematical biology: I. An introduction (Vol. 17). Springer Science & Business Media.

Nekouei, O., R. Vanderstichel, T. Ming, K. H. Kaukinen, K. Thakur, A. Tabata, E. Laurin, et al. 2018. Detection and Assessment of the Distribution of Infectious Agents in Juvenile Fraser River Sockeye Salmon, Canada, in 2012 and 2013. Frontiers in Microbiology 9.

Preikshot, D., R. J. Beamish, R. M. Sweeting, C. M. Neville, and T. D. Beacham. 2012. The Residence Time of Juvenile Fraser River Sockeye Salmon in the Strait of Georgia. Marine and Coastal Fisheries 4:438–449.

Purdon, A., M. A. Mole, M. J. Chase, and R. J. van Aarde. 2018. Partial migration in savanna elephant populations distributed across southern Africa. Scientific Reports 8:11331.

R Core Team. 2020. R: a language and environment for statistical computing. R Foundation for Statistical Computing, Vienna, Austria.

Rechisky E. L., A. D. Porter, S. D. Johnston, C. F. Stevenson, S. G. Hinch, B. P. V. Hunt, and D. W. Welch. 2021. Exposure Time of Wild, Juvenile Sockeye Salmon to Open-Net-Pen Atlantic Salmon Farms in British Columbia, Canada. North American Journal of Fisheries Management nafm.10574.

Ripple W. J., C. Wolf, T. M. Newsome, M. Galetti, M. Alamgir, E. Crist, M. I. Mahmoud, et al. 2017. World Scientists’ Warning to Humanity: A Second Notice. BioScience 67:1026–1028.

Runge C. A., T. G. Martin, H. P. Possingham, S. G. Willis, and R. A. Fuller. 2014. Conserving mobile species. Frontiers in Ecology and the Environment 12:395–402.

Salama, N. K. G., and A. G. Murray. 2011. Farm size as a factor in hydrodynamic transmission of pathogens in aquaculture fish production. Aquacult Environ Interact 2:61–74.

Santos, Y., F. Pazos, J. L. Barja, S. R. M. Jones, and L. Madsen. 2019. Tenacibaculum maritimum, causal agent of tenacibaculosis in marine fish.

Satterfield D. A., J. C. Maerz, and S. Altizer. 2015. Loss of migratory behaviour increases infection risk for a butterfly host. Proceedings of the Royal Society B: Biological Sciences 282:20141734.

Shea, D., A. Bateman, S. Li, A. Tabata, A. Schulze, G. Mordecai, L. Ogston, et al. 2020. Environmental DNA from multiple pathogens is elevated near active Atlantic salmon farms. Proceedings of the Royal Society B: Biological Sciences 287:20202010.

Soetaert, K., and F. Meysman. 2010. ReacTran: Reactive transport modelling in 1D, 2D and 3D. R package version 1.

Stevenson C. F., S. G. Hinch, A. D. Porter, E. L. Rechisky, D. W. Welch, S. J. Healy, A. G. Lotto, et al. 2019. The Influence of Smolt Age on Freshwater and Early Marine Behavior and Survival of Migrating Juvenile Sockeye Salmon. Transactions of the American Fisheries Society 148:636–651.

Taranger G. L., Ø. Karlsen, R. J. Bannister, K. A. Glover, V. Husa, E. Karlsbakk, B. O. Kvamme, et al. 2015. Risk assessment of the environmental impact of Norwegian Atlantic salmon farming. ICES Journal of Marine Science 72:997–1021.

Thakur K. K., R. Vanderstichel, S. Li, E. Laurin, S. Tucker, C. Neville, A. Tabata, et al. 2018. A comparison of infectious agents between hatchery-enhanced and wild out-migrating juvenile chinook salmon (*Oncorhynchus tshawytscha*) from Cowichan River, British Columbia. (S. J. Cooke, ed.)FACETS 3:695–721.

Tucker, S., M. Trudel, D. W. Welch, J. R. Candy, J. F. T. Morris, M. E. Thiess, C. Wallace, et al. 2009. Seasonal Stock-Specific Migrations of Juvenile Sockeye Salmon along the West Coast of North America: Implications for Growth. Transactions of the American Fisheries Society 138:1458–1480.

Vollset K. W., R. I. Krontveit, P. A. Jansen, B. Finstad, B. T. Barlaup, O. T. Skilbrei, M. Krkošek, et al. 2016. Impacts of parasites on marine survival of Atlantic salmon: a meta-analysis. Fish and Fisheries 17:714–730.

Welch D. W., M. C. Melnychuk, J. C. Payne, E. L. Rechisky, A. D. Porter, G. D. Jackson, B. R. Ward, et al. 2011. In situ measurement of coastal ocean movements and survival of juvenile Pacific salmon. Proceedings of the National Academy of Sciences 108:8708–8713.

Wiethoelter A. K., D. Beltrán-Alcrudo, R. Kock, and S. M. Mor. 2015. Global trends in infectious diseases at the wildlife–livestock interface. Proceedings of the National Academy of Sciences 112:9662–9667.

Wood C. C., J. W. Bickham, R. John Nelson, C. J. Foote, and J. C. Patton. 2008. Recurrent evolution of life history ecotypes in sockeye salmon: implications for conservation and future evolution: Recurrent evolution of ecotypes in sockeye salmon. Evolutionary Applications 1:207–221.

